# Comprehensive analysis of 111 Pleuronectiformes mitochondrial genomes: insights into structure, conservation, variation and evolution

**DOI:** 10.1101/2024.09.22.614327

**Authors:** Suxu Tan, Wenwen Wang, Jinjiang Li, Zhenxia Sha

## Abstract

Pleuronectiformes, also known as flatfish, are important model and economic animals. However, a comprehensive genome survey of their important organelles, mitochondria, is limited. In this study, we analyzed the genomic structure, codon preference, nucleotide diversity, selective pressure and repeat sequences, as well as reconstructed the phylogenetic relationship using the mitochondrial genomes of 111 flatfish species. Our analysis revealed a conserved gene content of protein-coding genes and rRNA genes, but varying numbers of tRNA genes across species. The mitochondrial genomes of most flatfish were conservative, while obvious gene rearrangements were found in several species, especially for the whole region rearrangement of *nad5*-*nad6*-*cytb* in Samaridae family and the swapping rearrangement of *nad6* and *cytb* gene in Bothidae family, suggesting a unique evolutionary history or functional benefit. Codon usage showed obvious biases, with adenine being the most frequent nucleotide at the third codon position. Nucleotide diversity and selective pressure analysis suggested that different protein-coding genes underwent varying degrees of evolutionary pressure, with *cytb* and *cox* genes being the most conserved ones. Phylogenetic analysis using both whole mitogenome information and concatenated independently aligned protein-coding genes largely mirrored the taxonomic classification of the species, with Samaridae family forming a distinct outgroup when the first approach was used. The identification of simple sequence repeats and various long repetitive sequences (forward, reverse, palindromic and complementary repeats) provided additional complexity of genome organization and offered markers for evolutionary studies and breeding practices. In summary, this study represents a significant step forward in our comprehension of the flatfish mitochondrial genomes, providing valuable insights into the structure, conservation and variation within flatfish mitogenomes, with implications for understanding their evolutionary history, functional genomics and fisheries management. Future research can delve deeper into the conservation biology, evolutionary biology and functional usages of variations.

## 1. Introduction

Fish are one of the most diverse and abundant vertebrate groups on Earth, playing crucial roles in aquatic ecosystems and global fisheries, as well as serving as important model organisms for scientific research (Nelson et al., 2016). Among them, the Pleuronectiformes, commonly known as flatfish, are particularly intriguing due to their unique morphology, ecological adaptations and economic significance (Munroe, 2014). These fish are characterized by their flattened bodies and asymmetric eyes, with both eyes shifted to one side of the head during the juvenile stage. This distinct morphology allows them to lie on the bottom of aquatic environments and ambush prey. The flatfish encompass a wide range of species distributed globally in marine, freshwater and brackish habitats, from shallow waters to deep-sea environments (Kenneth W. Able, 2014), contributing significantly to developmental and evolutionary studies. In addition, some flatfishes contribute to blue food provision and economic development, including flounders, soles and halibuts (FAO, 2024).

As the primary energy-generating system, mitochondria are essential organelles and found in virtually all eukaryotic cells. More than just powerhouses, they participate in multiple cell signaling cascades and perform a large suite of functions, including but not limited to development, metabolism, aging, disease and immunity (Chan, 2006; McBride et al., 2006; Mills et al., 2017). Mitochondria originated from a symbiotic relationship between a host prokaryote and an alpha-proteobacterium, therefore, they contain their own genome distinct from nuclear DNA although some genes were transferred to the host’s nuclear genome or lost during evolution (Andersson et al., 2003; Roger et al., 2017). The mitochondrial genome (mitogenome) is a circular DNA molecule with a relatively small size, typically ranging from 15 to 20 kilobases in vertebrate. Due to its maternal inheritance, lack of recombination and relatively fast evolutionary rate, mitogenome has become a powerful tool in phylogenetics and population genetics studies (Brown, 2008). In particular, complete mitogenome sequences offer a comprehensive view of genome rearrangement, codon usage and genetic diversity, thereby providing valuable insights into the evolutionary history of species. Additionally, analysis on mitogenome facilitates functional studies, and functional mitogenomic variations have been manifested by both laboratory experiments and disease phenotype. For instance, mutations in mitogenomes are associated with metabolic and neurological diseases in human (Ng and Turnbull, 2016). Using androgenesis rainbow trout (*Oncorhynchus mykiss*) with identical nuclear backgrounds and different maternal backgrounds, researchers revealed that mitochondrial variation could exert effects on growth rates and oxygen consumption, which may help increase food conversion ratios in aquaculture (Brown et al., 2006).

The comparative analysis of mitogenomes within the flatfish remains limited. Previous studies have focused primarily on individual genes or small genomic regions, such as cytochrome oxidase subunit I and 16S rRNA (Tang, 2011; Yang, 2010), for phylogenetic reconstruction and population genetic analyses. However, these approaches often suffer from limited information content and potential homoplasy issues (Iwasaki et al., 2013; Miya and Nishida, 2000). In contrast, the utilization of complete mitogenome sequences has the potential to overcome these limitations by providing a more holistic and comprehensive dataset. For instance, Shi (Shi, 2011) revealed mitogenome rearrangements, inversions and variations in a limited number of flatfish. Using complete mitogenome sequences of 39 flatfish species and other outgroup species, Campbell investigate the phylogenetic relationships within flatfish and provided weak support for the monophyly of flatfish (Campbell et al., 2014). Despite these advances, a comprehensive comparative and phylogenetic analysis encompassing a wide range of flatfish species is needed. In particular, repeat analyses are underexplored in flatfish, which would provide valuable resource for structural and functional implications. Such comprehensive analyses would not only fill gaps in our knowledge about the diversity of mitogenome within this fish group but also shed light on evolutionary history and functional significance.

The present study aims to bridge this gap by conducting a comparative and phylogenetic analysis of mitogenomes in 111 flatfish species. First, we seek to characterize the mitogenome features of the selected flatfish species, particularly in terms of gene content, gene rearrangement, codon usage, nucleotide diversity and selective pressure. Second, we intend to construct a robust phylogenetic framework for flatfish based on complete mitogenome sequences, providing new insights into the relationships among flatfish families and genera. Finally, by analyzing tandem repeats and dispersed repeats, we hope to provide valuable resources for better understanding of their functional implications and for potential applications in the selection or breeding of flatfish.

## 2. Material and Methods

### 2.1. Sequence retrieval

The complete mitogenome sequences of 111 flatfish species from 12 families were retrieved from the NCBI GenBank database (https://www.ncbi.nlm.nih.gov/). The species were chosen to cover a broad taxonomic range within the order Pleuronectiformes. The downloaded files included the gene sequences in FASTA format and their corresponding annotation information in GFF3 format.

### 2.2. Similarity comparison

To visually compare the mitogenomes of the 111 flatfish species, we used BLAST+ (Camacho et al., 2009) and BRIG (BLAST Ring Image Generator) (Alikhan et al., 2011). BRIG generates a circular visualization of multiple genomes, where the similarity between a reference genome and query sequences is shown as concentric rings based on BLAST consistency.

We grouped the 111 species into 6 sets, with *Paralichthys olivaceus* serving as the reference genome in each set. The first inner ring represented GC content, which is the percentage of guanine (G) and cytosine (C) nucleotides in the genome. The second inner ring depicted GC skew, calculated as the difference in the proportions of G and C in a sliding window, to identify functional regions potentially associated with replication (Sahyoun et al., 2014) and gene expression regulation (Hartono et al., 2015).

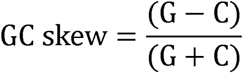

### 2.3. Collinearity analysis

Collinearity analysis was performed using Mauve software (Darling et al., 2004) to identify conserved and rearranged genomic regions among the 111 flatfish mitogenomes. The species were divided into 11 groups, and the Progressive Mauve algorithm was employed for multiple sequence alignment. Mauve connects similar sequence regions in different genomes with lines, representing conserved blocks. Nonlinear arrangements indicate genomic rearrangements such as insertions, deletions, or inversions.

### 2.4. Codon usage bias analysis

Codon usage bias, the preferential use of specific synonymous codons for the same amino acid, was analyzed. PhyloSuite (Zhang et al., 2020) was employed to extract the coding sequences of the protein-coding genes (PCGs) from the mitogenomes. Subsequently, codon usage indices, including CAI (Codon Adaptation Index), FOP (Frequency of Optimal Codons), CBI (Codon Bias Index), ENC (Effective Number of Codons) and GC3s (GC content at the third codon position), were calculated using CodonW (Peden, 2000). Using Enc and GC3s, Enc-plot was also conducted to evaluate the degree of preference for imbalanced use of synonymous codons. In addition, Relative Synonymous Codon Usage (RSCU) values were computed to identify codons with biased usage, where RSCU > 1 indicates that the frequency of use of the codon is higher than the average (positive bias), and RSCU < 1 indicates negative bias.

### 2.5. Nucleotide diversity analysis

The CDS sequences of the 13 PCGs were aligned using MAFFT in PhyloSuite, trimmed using Gblocks (Castresana and evolution, 2000), and concatenated into a single alignment file. Nucleotide diversity (Pi) was analyzed using DnaSP (Rozas et al., 2017), calculated using a sliding window approach with a window size of 200 and a step size of 20. The calculation formula for Pi value is as follows,

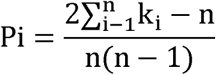

where N is the number of sequences in the sample, and k_i_ is the number of different nucleotides at the i^th^ site in the sequence.

### 2.6. Non-synonymous and synonymous substitutions

Evaluating sequence variations and evolution can be effectively achieved through calculating non-synonymous (Ka) and synonymous (Ks) substitution rates for protein orthologs. MUSCLE-generated (Edgar, 2004) protein alignments across species for each PCG were converted into codon comparisons using ParaAT v2.0 (Zhang et al., 2012), which were used for Ka/Ks calculation with KaKs_Calculator v2.0 (Wang et al., 2010).

### 2.7. Phylogenetic analysis

The complete mitogenome sequences of the 111 flatfish species were aligned using MAFFT in PhyloSuite. PCGs were first translated into amino acids, aligned, and then back-translated into nucleotides. The aligned sequences were optimized using MACSE (Ranwez et al., 2011), and trimmed using Gblocks selecting “Codons” as the data type. In contrast, RNA genes (tRNAs and rRNAs) were trimmed by Gblocks using “Nucleotide” as the data type after the alignment with MAFFT. All alignments were then concatenated together. PartitionFinder2 (Lanfear et al., 2017) was subsequently used to find the best partitioning strategy and to calculate the best-fit evolutionary models. A maximum likelihood phylogenetic tree was constructed using IQ-TREE (Nguyen et al., 2015), with SH-aLRT (Approximate Likelihood Ratio Test) to evaluate the reliability of the tree. The resulting tree was uploaded to iTOL (Letunic and Bork, 2024), and gene order was added for each mitogenome. In addition, the phylogenetic analysis was also performed based on 13 PCGs, where each of them was aligned separately and concatenated together, followed by the above-mentioned procedure.

### 2.8. Identification of simple sequence repeats (SSRs) and dispersed repeats

SSRs, also called microsatellites, were identified using MISA (MIcroSAtellite identification tool) (Beier et al., 2017), which scans the genome for repeated nucleotide motifs of different lengths. We set thresholds to detect SSRs of mono-, di-, tri-, tetra-, penta- and hexanucleotide repeats with minimum repeat units of 10, 5, 4, 3, 3 and 3, respectively. A minimum distance of 100 nucleotides was maintained between two SSRs to avoid counting overlapping regions.

Dispersed repeat sequences were analyzed using REPuter (Kurtz et al., 2001), which identifies forward, palindromic, reverse and complementary repeats. The minimum repeat size was set to 8 and the Hamming distance was 3 allowing for up to three mismatches. E-value of 0.01 was set as the threshold of significance.

## 3. Results

### 3.1. General features of flatfish mitogenomes

The mitogenome length, GC content percentage and NCBI accession number are listed in Table 1 for each studied species. All mitogenomes contain 13 PCGs (*atp6*, *atp8*, *cox1*, *cox2*, *cox3*, *cytb*, *nad1*, *nad2*, *nad3*, *nad4*, *nad4L*, *nad5* and *nad6*) and two rRNA genes (12S rRNA and 16S rRNA), but have different numbers of tRNA genes. Specifically, not all species have 22 tRNA genes, with a set of 24 species harboring 20 tRNAs and five harboring 21 tRNAs. The distribution of gene length, GC content and GC skew for all species are shown in Fig. 1. The gene length for each gene is similar across studies species, with very small variations (Fig. 1A). This is also true for two rRNA genes, with the length of 12S rRNA genes being approximately 950 bp and 16S rRNA genes being around 1700 bp, and the wide distribution in the figure is solely because they are plotted together. Different genes have varying GC content, and most of them are below 50%, consistent with the overall GC percent for mitogenome. Compared with GC content, the GC skew of different genes exhibit more variations. The *apt8* gene shows the lowest GC skew, and the *nad6* gene shows the highest GC skew, which is also the only PCG with positive GC skew value. Generally, the rRNA and tRNA genes exhibit higher GC skew than PCGs. The GC and GC skew of tRNA genes exhibit the greatest degree of variability, which may be due to the large number of genes plotted together.

**Fig. 1.**
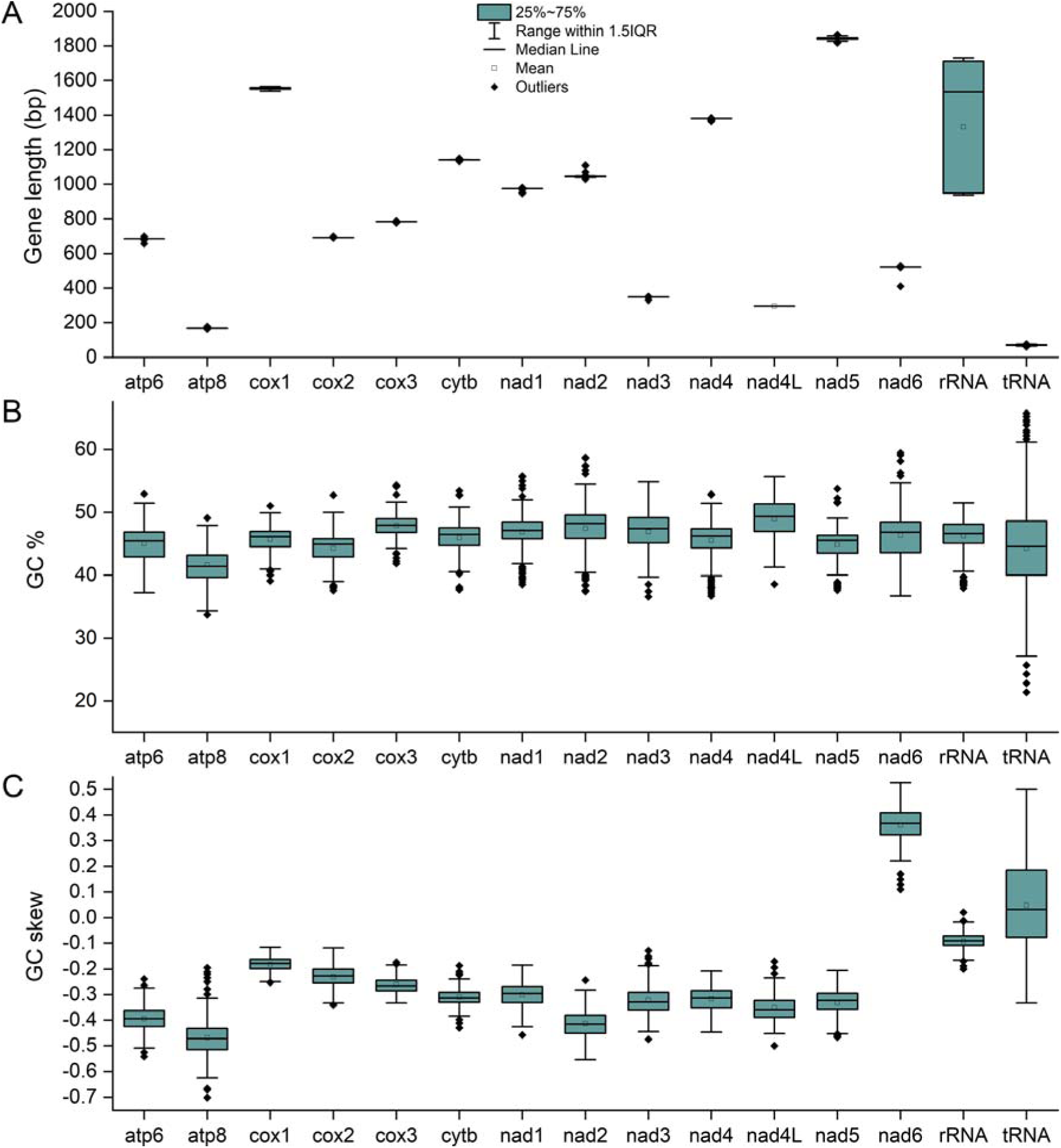
The distribution of gene length (a), GC base composition (b) and GC skew (c) in flatfish mitogenomes.

**Table 1.**
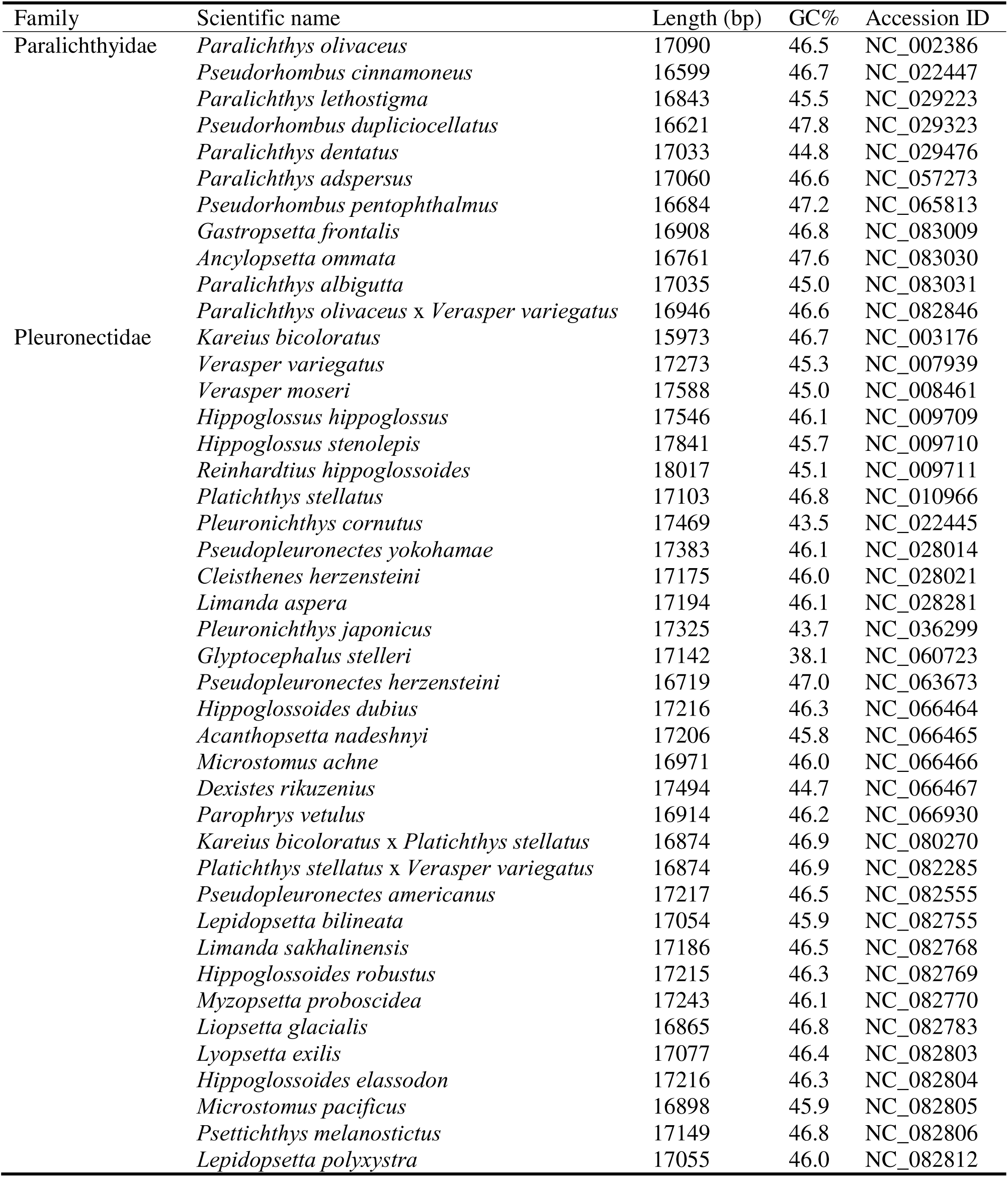
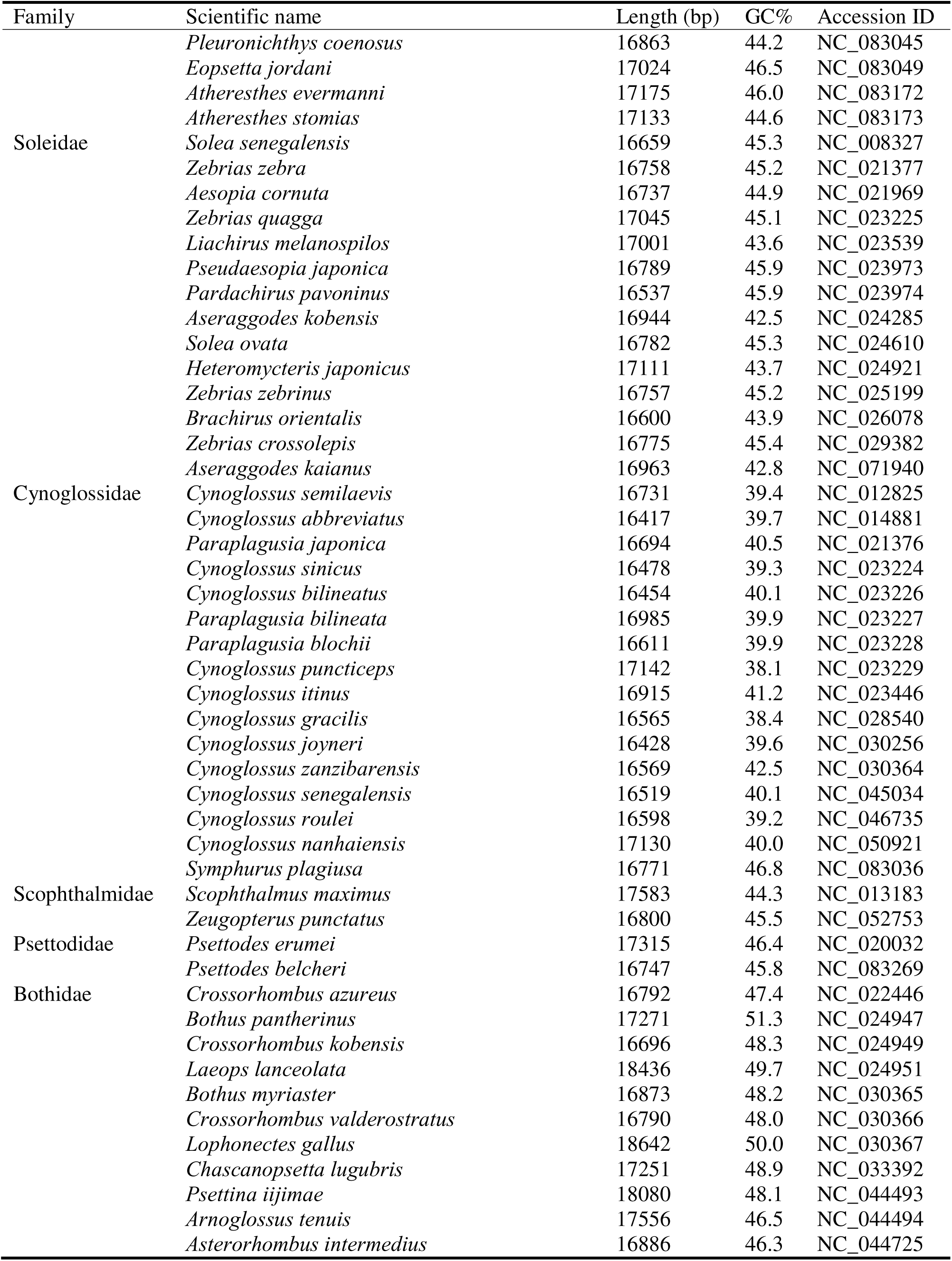
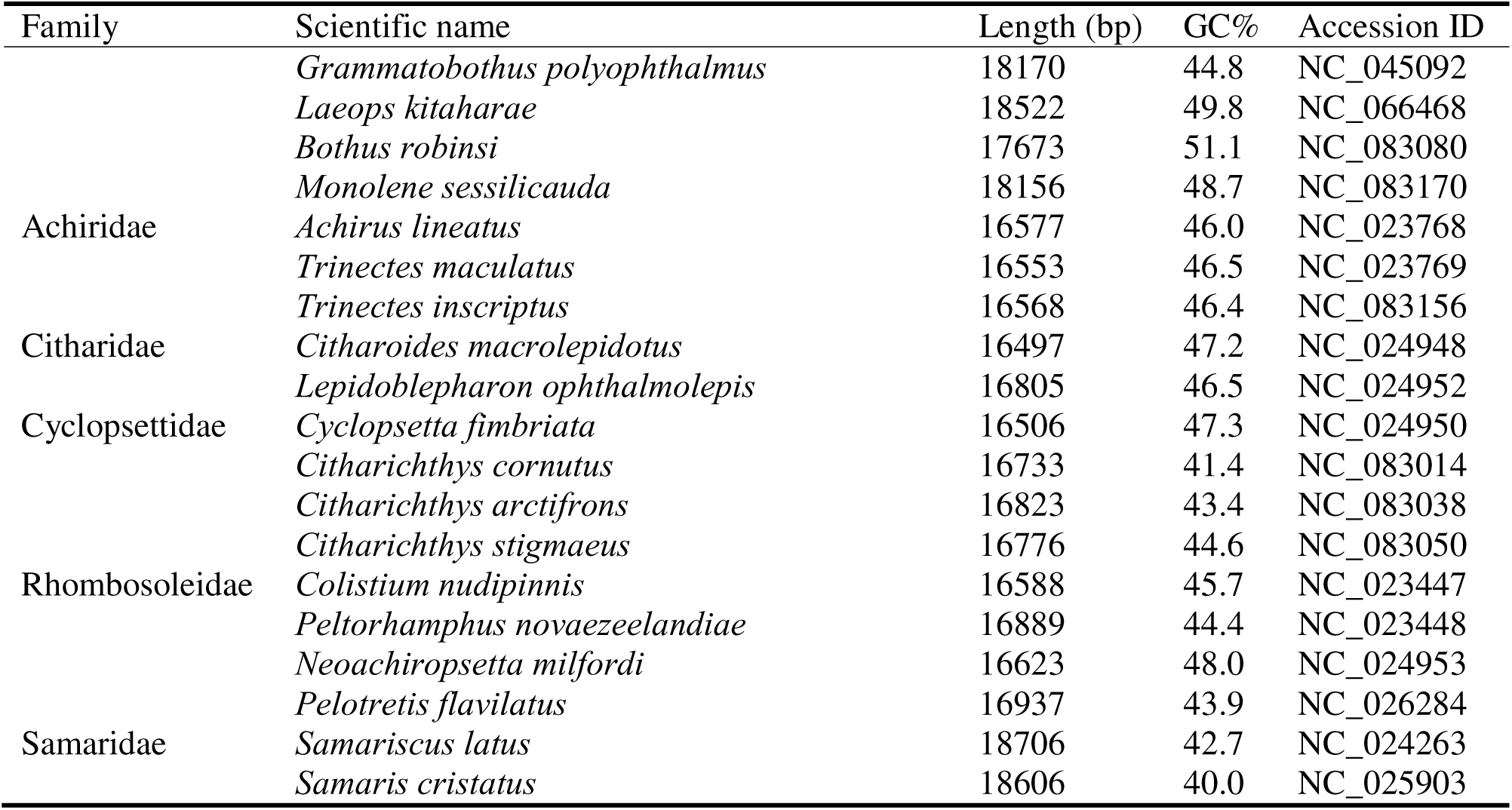
Species information and genome characteristics.

### 3.2. Similarity among flatfish mitogenomes

The mitogenomes of the 111 Pleuronectiformes species were compared to assess their similarity. The alignment of 20 representative species (designated Group 1) are shown in Fig. 2, and the other five groups are shown in Fig. S1. The results indicate that mitogenomes exhibited high similarity with most sequences exhibiting over 70% similarity, indicating a high level of homology among the genomes. GC content and GC skew appear randomly distributed throughout the majority of the genome. However, variations and peaks exist, which may have implications for their mitochondrial function and evolution, except for the peaks in the unaligned breaks in the upper area of the image, where biased GC skew only corresponds to the selected reference mitogenome of *P. olivaceus*.

**Fig. 2.**
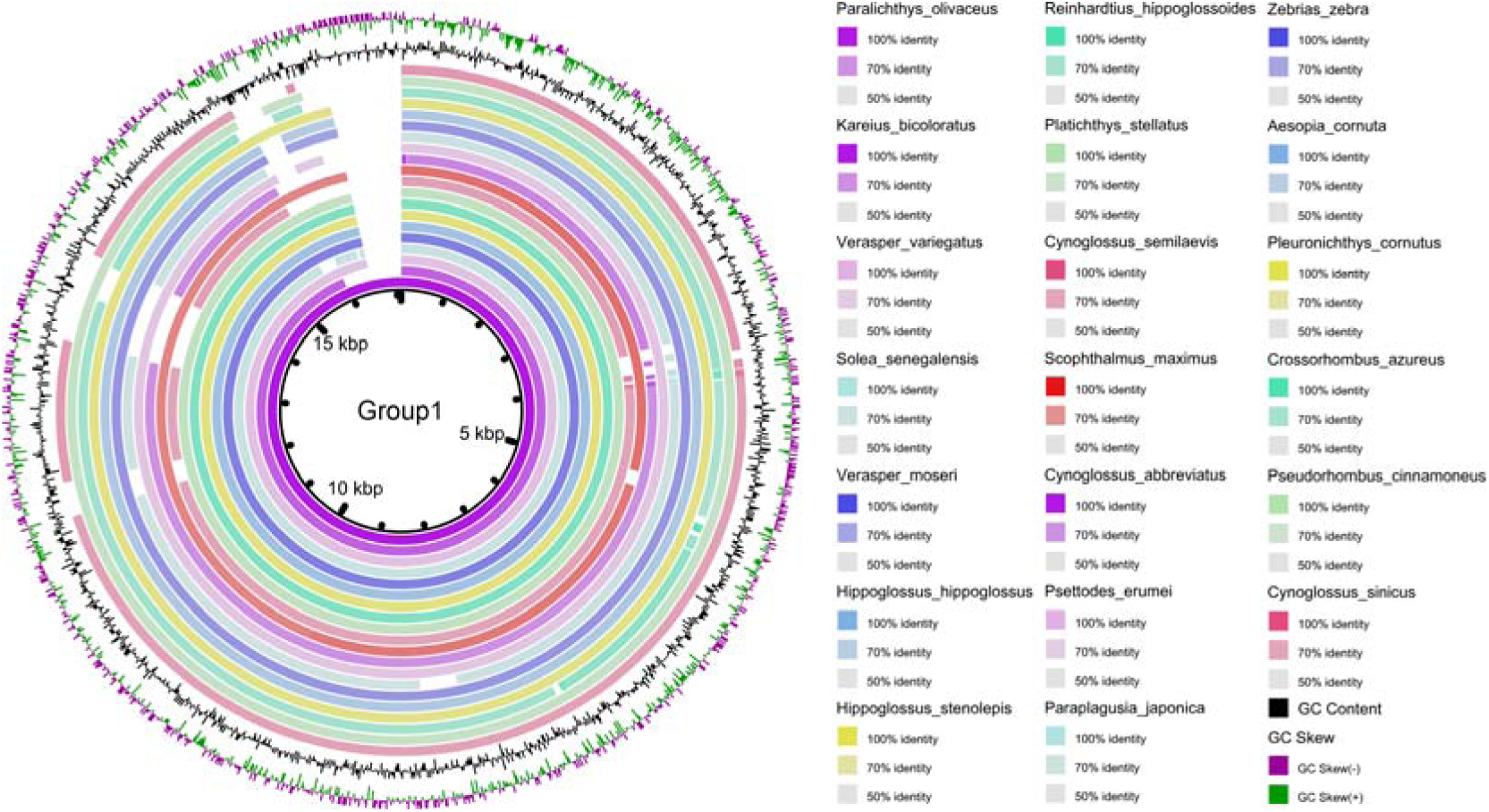
Alignment of mitochondrial genomes among studied representative Pleuronectiformes species. The gap in the circle represents mismatched sequence of genome alignment. GC content (black) and GC skew (purple/green) are at the outermost two circles. Increased GC content and positive GC skew are represented by peaks oriented toward the center of the circle.

### 3.3. Genomic rearrangement

To reveal the conservation among mitogenomes, the collinearity analysis was performed. Figure 3 shows the collinearity of mitogenomes of 24 representative species, including two species from each family. Comprehensive illustration of 111 species is shown in Fig. S2. Collinearity analysis showed strong conservation in gene arrangement of flatfish, with numerous homologous co-linear blocks being observed. There are also many exceptions. Specifically, *Samariscus latus* and *Samaris cristatus* exhibited the most significant genomic rearrangements, i.e., the green block (including *nad5*-*nad6*-*cytb* region) transfer from the right side of the light green block to the left side. They both belong to the Samaridae family (Table 1), and this family only contained these two species in the study. Moreover, a total of 15 species exhibited swapping gene rearrangement between *nad6* and *cytb* (Fig. S2); they all belong to the Bothidae family (Table 1), and this family contained these 15 species in the study. These findings indicated that the genomic region near genes of *nad5*, *nad6* and *cytb* might be a hotspot for rearrangement events during the evolution of flatfish mitogenomes. These results also suggested the unique evolution status of these two families and the functional implication of the genomic rearrangement. In addition, the dark red co-linear block in non-protein coding region exhibits high variations across the flatfish mitogenomes, indicating highly unconserved structure. In addition, certain regions (gaps between colored blocks) exhibited no homology, demonstrating their exclusive presence in this mitogenome.

**Fig. 3.**
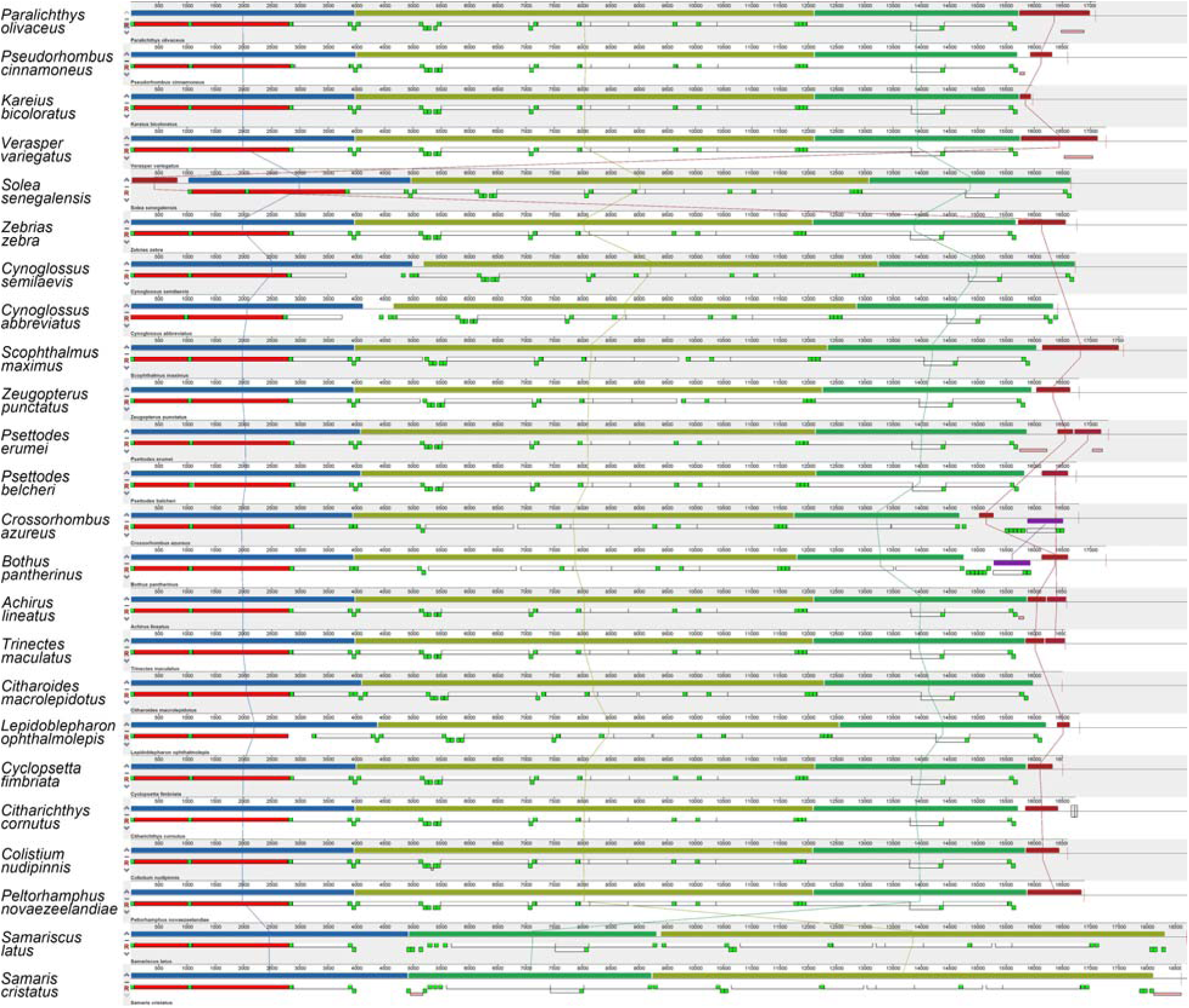
Collinearity analysis of mitochondrial genomes in part of studied flatfish, two from each family. For each species, the first row represents the collinear blocks, and the second row represents the genes, with rRNA genes indicated in red, protein-coding genes in white, tRNA genes in green, and repeat region in pink. co-linear blocks are connected by lines of the same color.

### 3.4. Codon preference of PCGs

To gain insights into the gene expression regulation and evolution, codon usage analysis was performed. The analysis revealed significant biases in the usage of specific codons among the flatfish mitogenomes. The top five frequently used codons are CUA, AUU, CUU, CUC, GCC, corresponding to the amino acids leucine (Leu), isoleucine (Ile), Leu, Leu and alanine (Ala), respectively (Table 2). In terms of RSCU, a total of 31 codons are positively biased, with CGA, CCC, GCC, UCA and CAA being the most significant ones, indicating a strong preference for these codons. Similarly, a set of 28 codons are negatively biased, with GCG, ACG, CCG, UUG, AGU being the most biased ones. The only codon without obvious usage preference is CCU, whose RSCU value is equal to one. The patterns of RSEU are consistent among the analyzed species (Fig. S3), with part of them showing in Fig. 4.

**Fig. 4.**
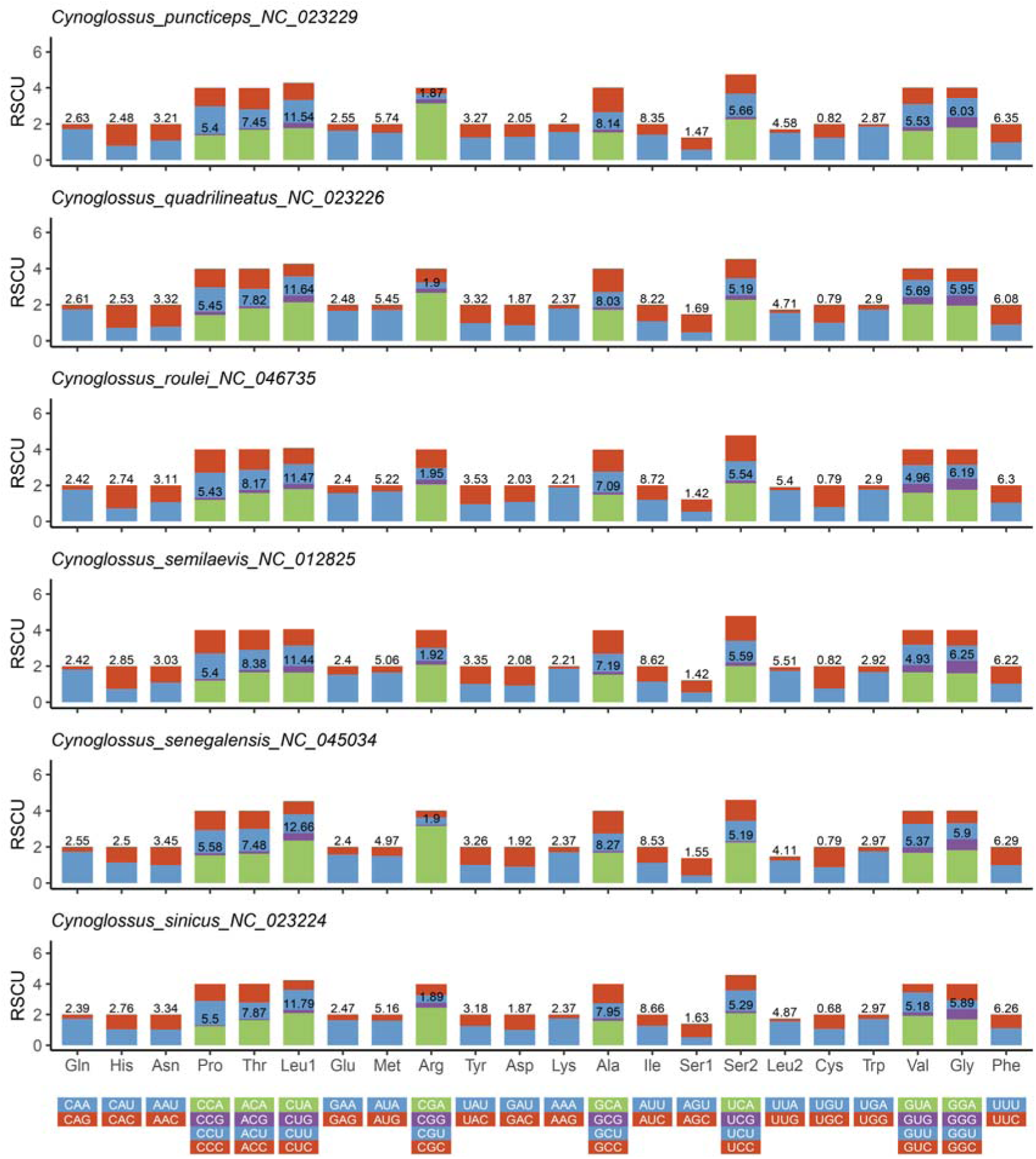
Relative synonymous codon usage of protein-coding genes in Pleuronectiformes mitogenomes

**Table 2.**
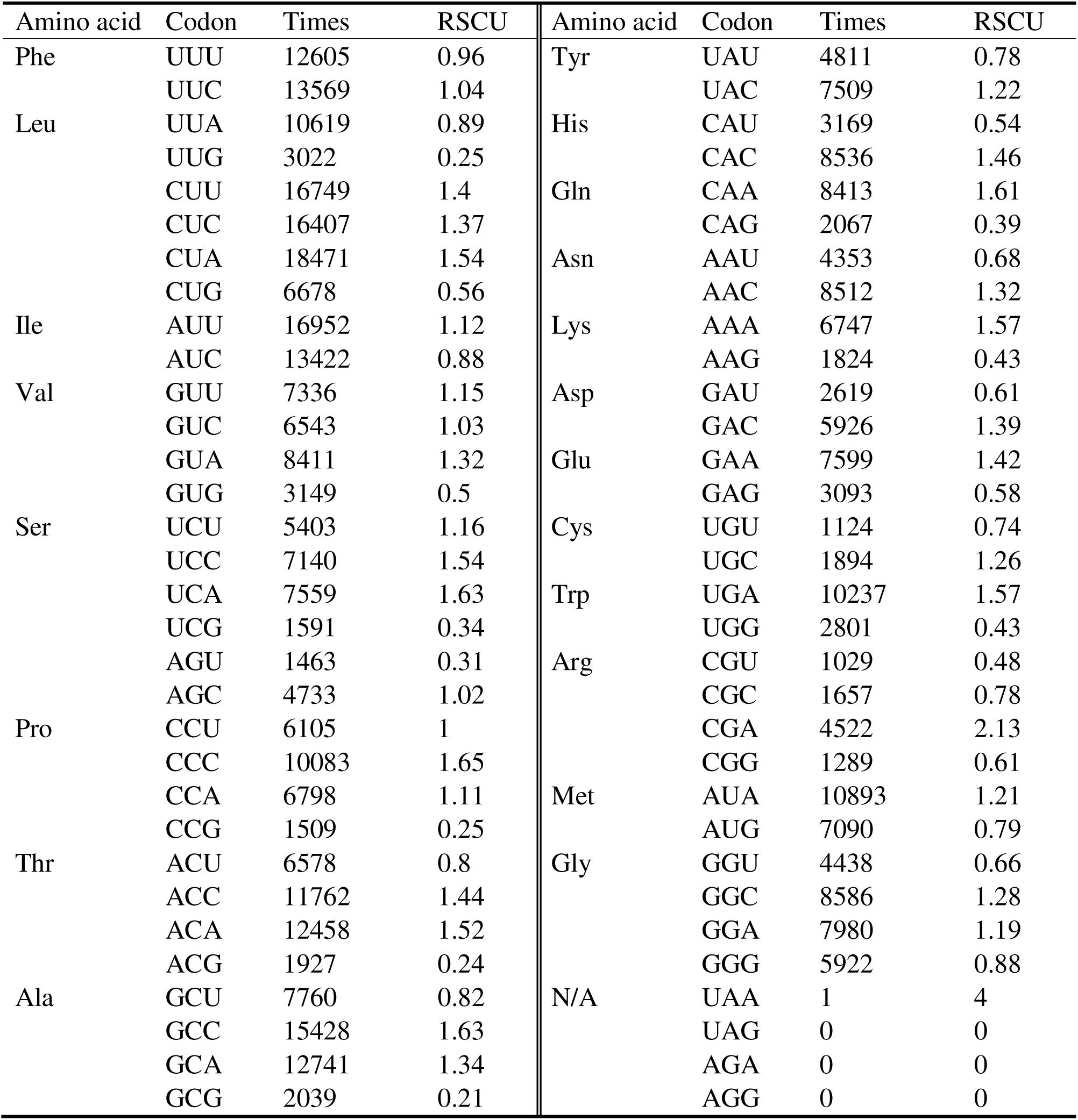
Preference of codon usage in the mitochondrial genomes of Pleuronectiformes.

Furthermore, a comparative analysis of codon preferences was conducted for each PCG (Table 3), demonstrating that codon bias varied between the genes. In addition, several characteristics were revealed. The frequency of adenine at the third position of synonymous codons (A3s) was the highest in most genes, accounting for 0.422 on average. This was in accordance with the RSCU results, i.e., most codons with RSCU > 1 were end in A (Table 2, Fig. 4). In contrast, guanine at the third position (G3s) had the lowest frequency of 0.144 on average. The GC content at the third position (GC3s) was 0.44, which was roughly equivalent to the overall GC content (0.46). The frequency of optimal codons (FOP) ranged from 0.33 to 0.43. Noting that FOP = 1 indicates that the gene exclusively uses optimal codons, while FOP = 0 suggests that the gene does not use any optimal codons. The codon adaptation index (CAI) value ranged from 0.134 to 0.187, indicating a relatively low degree of match between the codon usage patterns of these genes and those of highly expressed genes. It is noted that a CAI value closer to 1 indicates that the codon usage pattern of a gene is more akin to the preferences observed in highly expressed genes. The range of ENC values is 45.8 to 49.5 for genes in this study, and the overall Enc value is 20 to 61, where 20 indicates that only one codon is used for each amino acid, and 61 indicates that each codon is evenly used. The lower the ENC value, the stronger the preference for codon usage.

**Table 3.** Codon preference analysis of 13 protein-coding genes in Pleuronectiformes.

| Gene | T3s | C3s | A3s | G3s | CAI | CBI | FOP | ENC | GC3s |
| --- | --- | --- | --- | --- | --- | --- | --- | --- | --- |
| *atp6* | 0.298 | 0.364 | 0.421 | 0.116 | 0.134 | -0.080 | 0.33 | 48.0 | 0.41 |
| *atp8* | 0.299 | 0.402 | 0.494 | 0.125 | 0.150 | 0.016 | 0.41 | 47.5 | 0.40 |
| *cox1* | 0.298 | 0.383 | 0.436 | 0.142 | 0.174 | -0.033 | 0.40 | 48.0 | 0.43 |
| *cox2* | 0.310 | 0.394 | 0.480 | 0.121 | 0.187 | -0.019 | 0.41 | 47.4 | 0.41 |
| *cox3* | 0.270 | 0.438 | 0.447 | 0.101 | 0.175 | 0.008 | 0.43 | 46.3 | 0.45 |
| *cytb* | 0.264 | 0.447 | 0.421 | 0.114 | 0.179 | 0.021 | 0.42 | 47.4 | 0.48 |
| *nad1* | 0.272 | 0.392 | 0.414 | 0.139 | 0.154 | -0.044 | 0.37 | 49.1 | 0.45 |
| *nad2* | 0.266 | 0.414 | 0.401 | 0.121 | 0.144 | -0.047 | 0.37 | 48.5 | 0.45 |
| *nad3* | 0.280 | 0.410 | 0.409 | 0.135 | 0.141 | -0.055 | 0.35 | 46.9 | 0.45 |
| *nad4* | 0.270 | 0.385 | 0.436 | 0.137 | 0.137 | -0.033 | 0.38 | 49.3 | 0.43 |
| *nad4L* | 0.233 | 0.399 | 0.421 | 0.123 | 0.148 | -0.050 | 0.37 | 45.8 | 0.46 |
| *nad5* | 0.275 | 0.426 | 0.425 | 0.128 | 0.177 | 0.002 | 0.42 | 49.5 | 0.46 |
| *nad6* | 0.409 | 0.122 | 0.276 | 0.371 | 0.155 | -0.108 | 0.33 | 47.1 | 0.41 |
| Average | 0.288 | 0.383 | 0.422 | 0.144 | 0.158 | -0.032 | 0.38 | 47.7 | 0.44 |

To investigate the role of mutational pressure in determining the codon usage bias, the ENC-plot analysis was conducted. As shown in Fig. 5, the actual ENC values of all genes are below the expected curve, suggesting that mutation pressure is not the primary factor influencing codon usage and that natural selection plays a significant role.

**Fig. 5.**
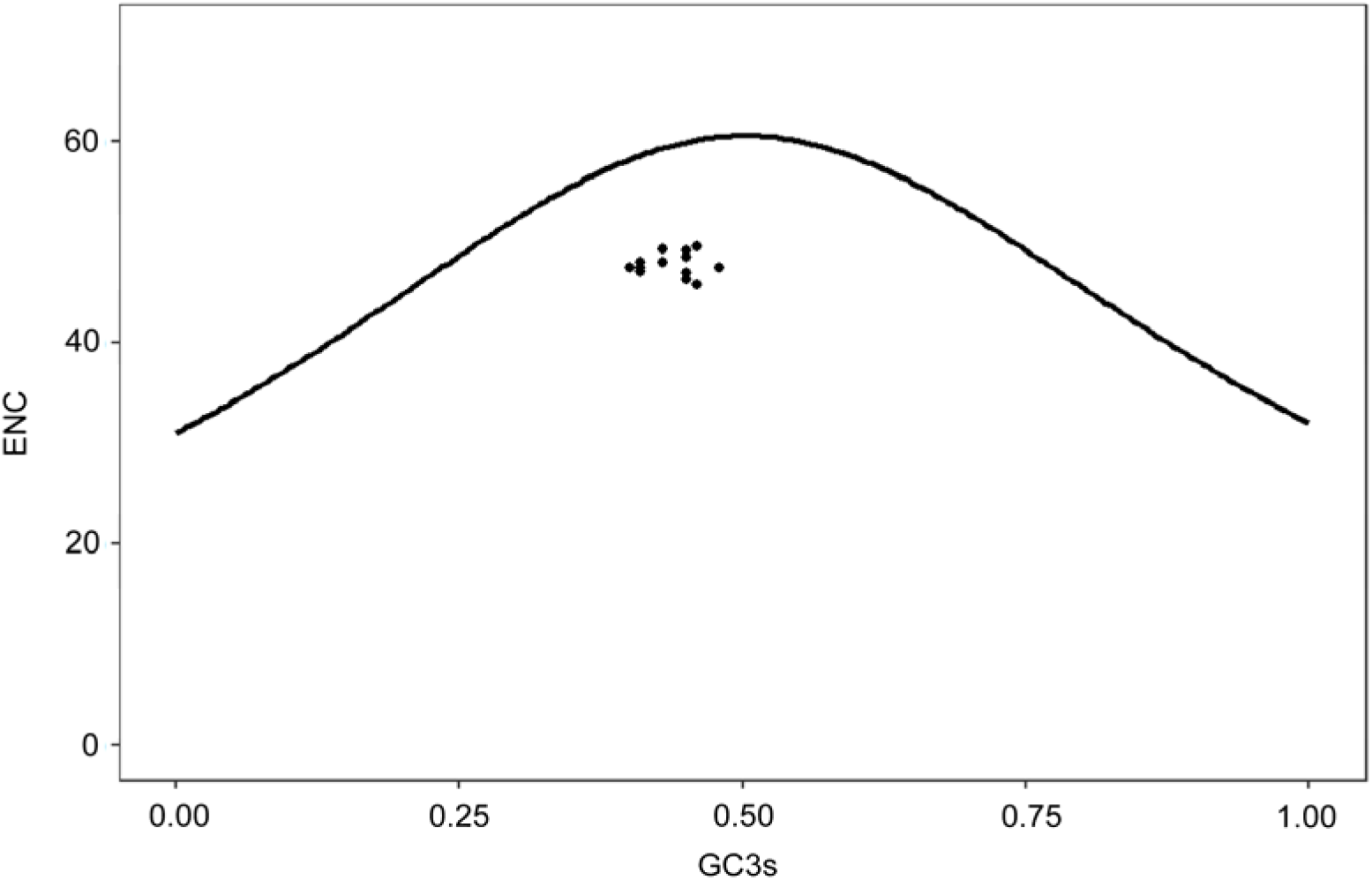
ENC-plot of codon usage in mitochondrial genome of Pleuronectiformes. GC3s, GC content at the third codon position; ENC, effective number of codons.

### 3.5. Nucleotide diversity of PCGs

Nucleotide diversity analysis of the 13 PCGs revealed varying levels of nucleotide diversity among the genes, ranging from 0.207 to 0.359 (Fig. 6). The *atp8* gene exhibited the highest nucleotide diversity (Pi = 0.359), followed by *nad6* (Pi = 0.327), *nad2* (Pi = 0.323) and *atp6* (Pi = 0.306). In contrast, *cox1* (Pi = 0.207), *cox2* (Pi = 0.249), *cox3* (Pi = 0.234) and *cytb* (Pi = 0.242) were relatively conserved with lower Pi values. These findings suggest that different genes may undergo distinct evolutionary pressures, leading to variations in their nucleotide diversity. Moreover, different regions of each gene showed varying levels of nucleotide diversity, providing valuable information for primer designing to distinguish species or conduct evolutionary analysis.

**Fig. 6.**
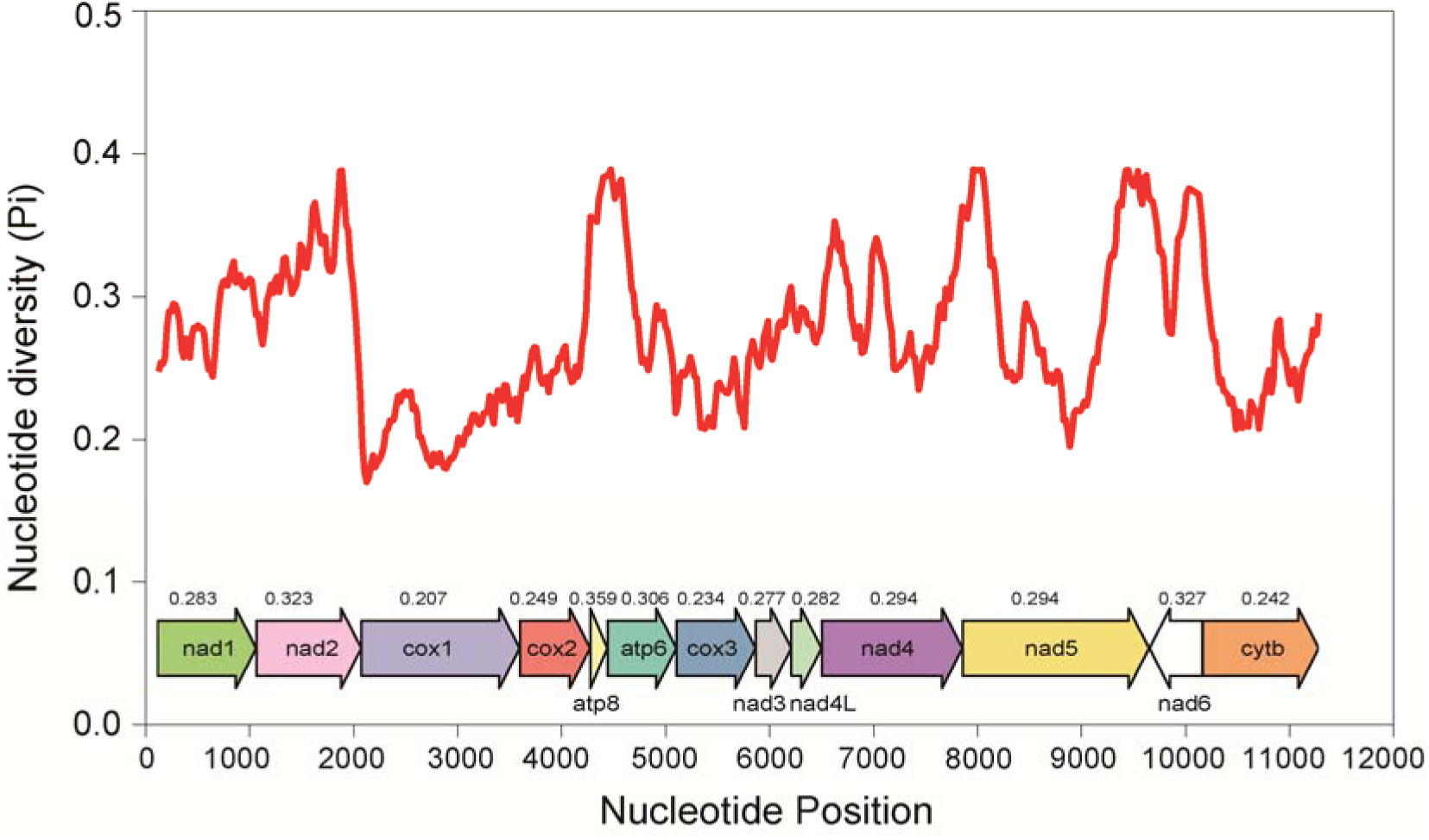
Nucleotide diversity in the 13 protein-coding genes in the mitochondrial genomes of Pleuronectiformes. The X axis denotes the nucleotide position of PCGs, and the Y axis denotes the nucleotide diversity (Pi) value. The continuous red line represents the nucleotide diversity value in different regions of the protein coding gene. The number above each gene represents the average Pi value of the gene.

### 3.6. Selective pressure of PCGs

To reveal the selective pressure of each mitochondrial PCG in flatfish, the Ka/Ks value was calculated. As shown in Fig. 7, purifying selection exerts great influence on all the PCGs. Specifically, the *atp8* gene shows the highest Ka/Ks value, and the cox family genes are generally low with *cox1* being the lowest. This pattern is consistent with the result of nucleotide diversity analysis, reinforcing the genetic conservation and evolutionary pressure of each PCG in flatfish mitogenome. Intriguingly, this phenomenon is widely found in most Metazoa (Castellana et al., 2011).

**Fig. 7.**
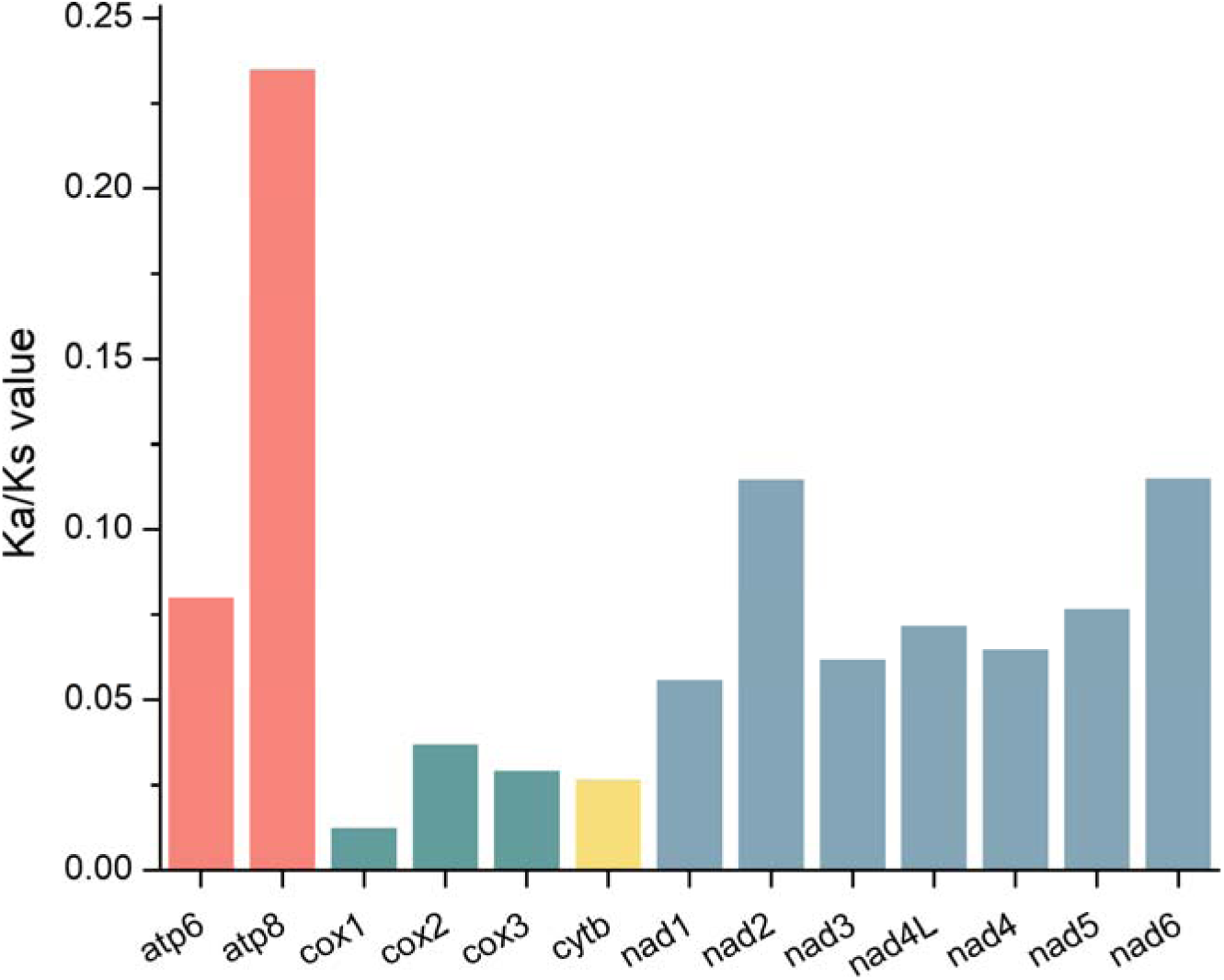
Non-synonymous/synonymous ratio (Ka/Ks) of the 13 protein-coding genes in flatfish mitogenomes. The genes (*apt6* and *atp8*) encoding ATPase complex are indicated in red, the genes (*cox*1-3) encoding cytochrome c oxidase are indicated in green, the gene (*cytb*) encoding cytochrome b is represented in orange, and the genes (*nad*1-6 and 4L) encoding NADH dehydrogenase complex are represented in blue.

### 3.7. Phylogenetic reconstruction

The phylogenetic analysis of the 111 Pleuronectiformes species revealed a well-resolved phylogenetic tree that largely mirrored the taxonomic classification of the species. The phylogenetic tree using the mitogenome (PCGs and RNA genes) revealed that species were grouped according to their families and genera, with high bootstrap support values indicating robust phylogenetic relationships (Fig. 8). The exceptions are *Atheresthes stomias* and *Atheresthes evermanni* (highlighted in red arrow) who belong to the family Pleuronectidae grouped to the clade of Paralichthyidae. Intriguingly, *S. latus* and *S. cristatus* deviated significantly from other species, consistent with the observed gene rearrangements in these two species.

**Fig. 8.**
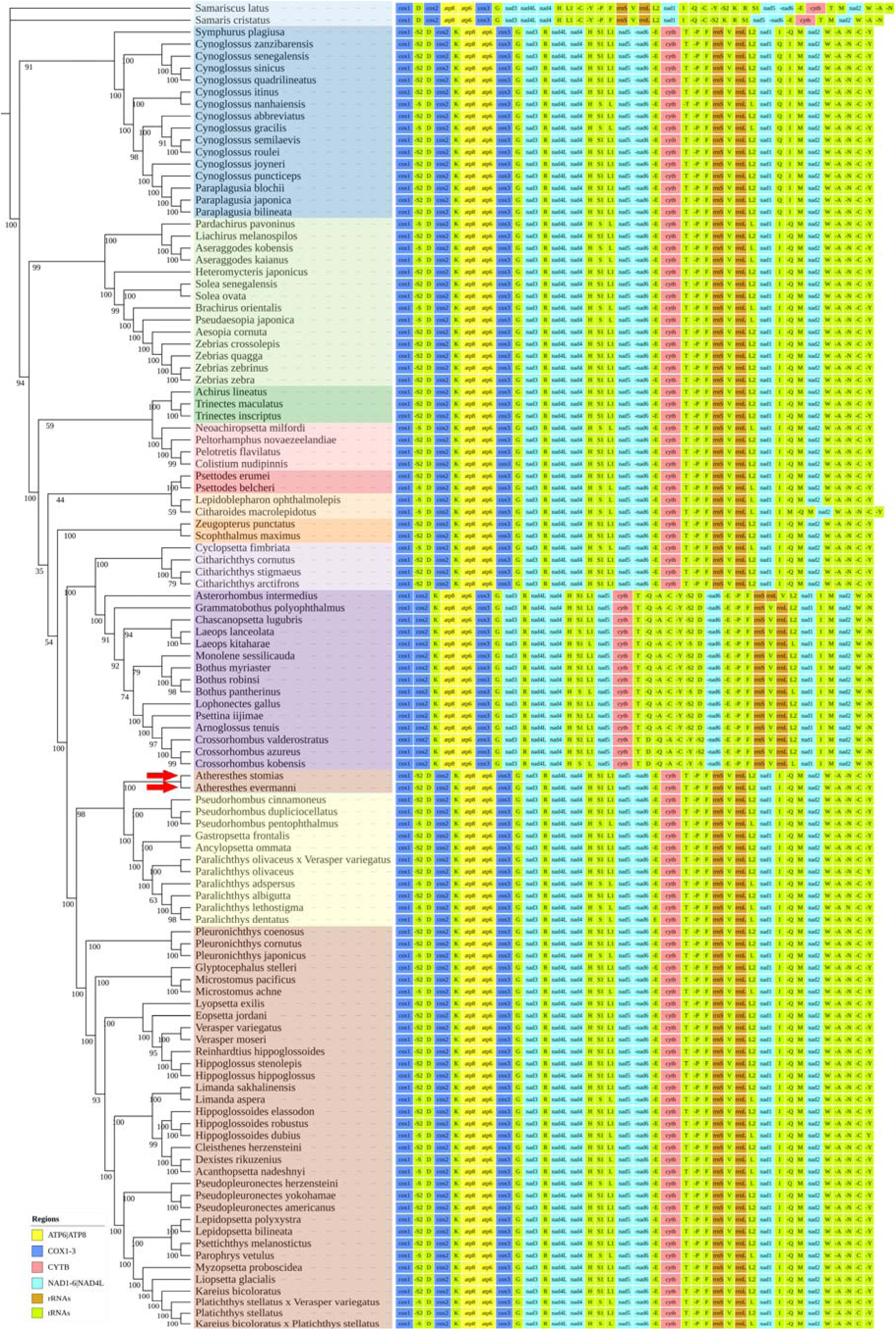
Phylogenetic tree constructed using whole mitogenome (PCGs and RNA genes) of studied Pleuronectiformes. In the left panel, different colors represent different families: light blue, Samaridae; blue, Cynoglossidae; light green, Soleidae; green, Achiridae; pink, Rhombosoleidae; red, Psettodidae; light orange, Citharidae; orange, Scophthalmidae; light purple, Cyclopsettidae; purple, Bothidae; yellow, Paralichthyidae; brown, Pleuronectidae. In the right panel, gene order for each mitogenome is provided, with a color legend displayed in the bottom left corner. The red arrows indicate species which are not grouped into their belonged family.

The phylogenetic tree constructed using concatenated separately aligned PCGs showed similar results that clades mostly correspond to the taxonomic classification (Fig. 9). Without the gene rearrangement information, the family Samaridae (*S. latus* and *S. cristatus*) does not form a distinct clade away from all other species. In addition, *Symphurus plagiusa* (highlighted in blue arrow) of family Cynoglossidae is classified into the family Samaridae in the tree, but the incongruity is not observed in Fig. 8. These findings provide new insights into the evolutionary history of Pleuronectiformes and support the utility of mitogenome sequences, instead of single gene, for phylogenetic reconstruction.

**Fig. 9.**
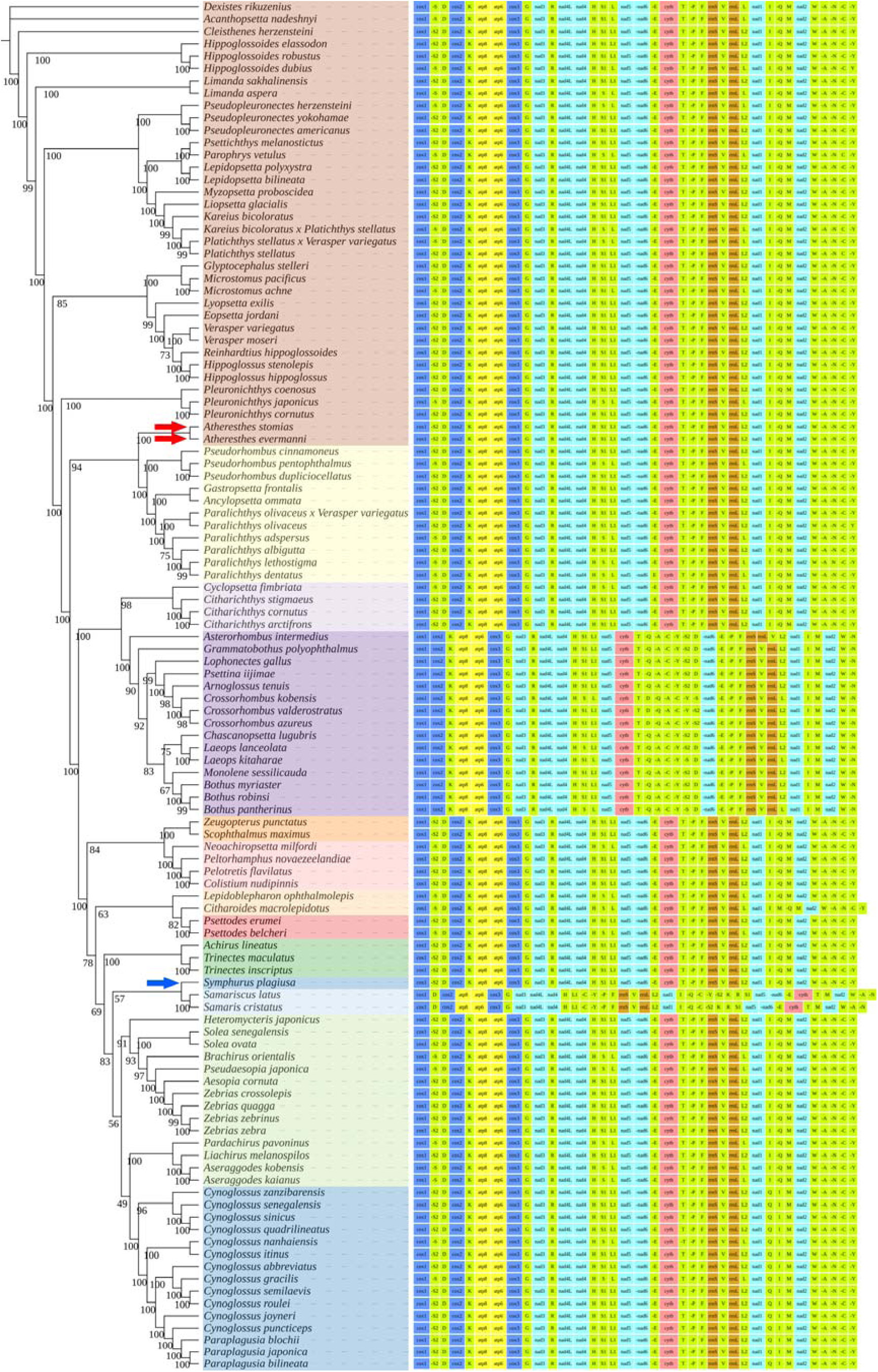
Phylogenetic tree constructed using PCGs of studied Pleuronectiformes mitogenomes. In the left panel, different colors represent different families: light blue, Samaridae; blue, Cynoglossidae; light green, Soleidae; green, Achiridae; pink, Rhombosoleidae; red, Psettodidae; light orange, Citharidae; orange, Scophthalmidae; light purple, Cyclopsettidae; purple, Bothidae; yellow, Paralichthyidae; brown, Pleuronectidae. In the right panel, gene order for each mitogenome is provided, with a color legend displayed in the bottom left corner. The red and blue arrows indicate species which are not grouped into their belonged family. In addition, the blue arrow also indicates the species whose classification is not consistent with that in Fig. 8.

### 3.8. SSR and dispersed repeat analysis

The SSRs identified in each species are shown in Table S1. Overall, a total of 227 SSR sites were identified across the studied mitogenomes. Most SSR types were monomeric (43.6%), followed by trimeric (18.9%) and tetrameric (14.1%). The most frequent monomeric SSR was an T repeat monomer, occurring 59 times and making up 59.6% of the total. The most common dimeric and trimeric SSR was AT/TA and TCC. It is noteworthy that mononucleotide A and hexanucleotide repeat were not identified in these mitogenomes. For each species, the number of total SSRs was from zero to seven, with 31 species containing one SSR, 40 species containing 2 SSRs and 21 species containing three SRRs. No SSR was identified in seven species, and *Bothus myriaster* harbored the most SSR sites, including five monomeric, one trimeric and one tetrameric site.

In addition to simple sequence repeat SSR, we also revealed the presence of forward, palindromic, reverse and complementary repeats in flatfish species. The forward, reverse, palindromic and complementary repeats measured 17-1178, 17-186, 17-41 and 17-38 bp, respectively (Table S2). In general, forward repeats were the most frequent, followed by reverse, palindromic and complementary repeats (Fig. 10 and Table S2). These findings suggest that SSRs and repeat sequences are prevalent in Pleuronectiformes mitogenomes, which may serve as useful molecular markers for evolutionary studies or breeding practices.

**Fig. 10.**
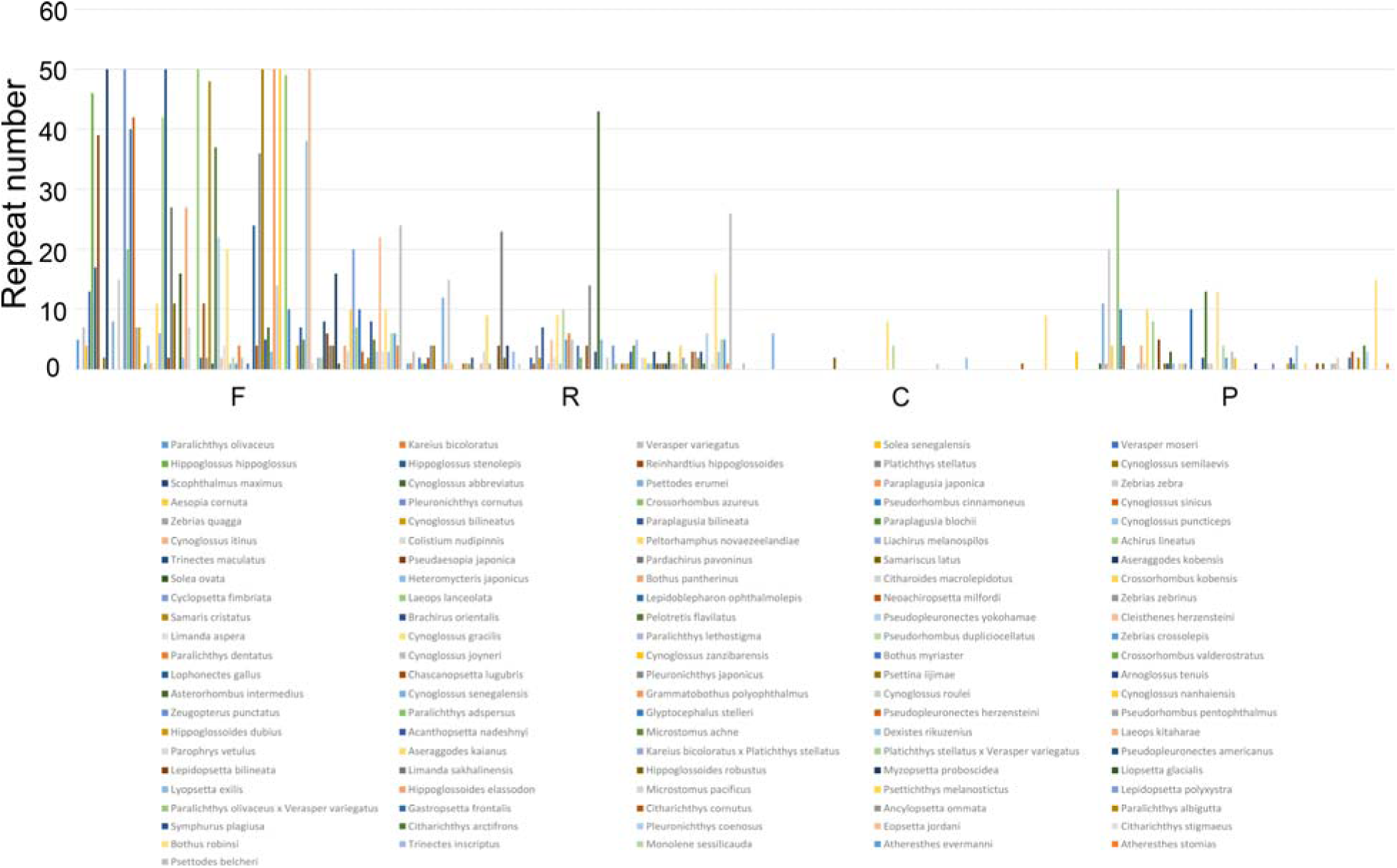
Numbers of four long repetitive sequences in the mitochondrial genomes of studied Pleuronectiformes. F, R, C, and P represent forward, reverse, complementary, and palindromic repeats, respectively. Species are indicated by different colors and annotations are displayed at the bottom of the image.

## 4. Discussion

With the advent of sequencing technology, a large number of mitogenomes have been sequenced, providing valuable resources for evolutionary, structural and functional analyses. The present study provides a comprehensive analysis of the mitogenomes of flatfish species, revealed several intriguing aspects of the structure, conservation and variation within these genomes. Our findings not only shed light on the evolutionary trajectories of these important economic and model organisms but also offer valuable insights into their functional genomics.

Our findings reveal a high degree of conservation in flatfish mitogenomes. The consistent gene length across flatfish species may be indicative of the conserved nature of mitochondrial genes. Though each PCG exhibited length variation, the variation in the *cox1* and *nad5* was found to be the largest in this study, whose gene size was also the largest. This result is consistent with previous findings (Satoh et al., 2016), which showed similar results among 250 fish species and suggested the close relationship between gene size and length variation. The observed variations in GC skew among different genes may be attributed to selective pressures or differences in transcriptional activity. The GC skews of PCGs were all negative except for *nad6*, which was also observed in other species, such as croakers and *Gerres* species (Ruan et al., 2020; Yang et al., 2018). The reason is that *nad6* is the only gene on the light strand (L-strand) and all other PCGs are located on the heavy strand (H-strand), which demonstrated the strand asymmetry and the patterns of nucleotide composition in flatfish mitogenome. The higher GC skew in rRNA and tRNA genes compared to PCGs could be related to their functional roles in protein synthesis and ribosome assembly. For example, GC skew at the 5′ and 3′ ends of human genes was correlated with R-loop formation which could affect epigenetic and transcription regulation (Ginno et al., 2013).

Synonymous codon usage bias is observed in all domains of life, and every organism possesses a unique codon choice affecting gene expression, translational efficiency, protein folding, protein function and therefore organismal fitness (Buhr et al., 2016; Quax et al., 2015; Yu et al., 2015). The codon usage analysis presented in this study provides valuable insights into the gene expression regulation and evolutionary dynamics of flatfish mitogenomes. The observed biases in codon usage suggest that natural selection has played a significant role in shaping the genetic code of these organisms. The predominance of codons ending in A at the third position of synonymous codons is consistent with the RSCU analysis, indicating a strong preference for these codons. This pattern is generally consistent across vertebrate groups, and this preference may be related to factors such as tRNA availability, translational efficiency, or codon-anticodon interactions (Broughton et al., 2001). The ENC-plot analysis indicates that natural selection is the primary determinant of codon preference in flatfish mitogenomes, while mutation only accounted for part of the reason (Wright, 1990). This suggests that codon usage is optimized for specific functional requirements. Further investigations into the underlying mechanisms driving codon bias and the functional implications of codon usage variations could provide valuable insights into the adaptive strategies of these organisms.

The results of collinearity analysis demonstrate a high degree of gene order conservation across the studied species, suggesting a shared evolutionary history. However, the presence of genomic rearrangements in certain species highlights the potential for lineage-specific evolutionary events, especially for the whole region rearrangement of *nad5*-*nad6*-*cytb* in Samaridae family and the swapping rearrangement of *nad6* and *cytb* gene in Bothidae family. The most striking example of genomic rearrangement is observed in the two species of the Samaridae family, *S. latus* and *S. cristatus*. The translocation of *nad5*-*nad6*-*cytb* gene block suggests a unique evolutionary history for this family. Gene rearrangements in animal mitogenomes are usually explained by three main models: the recombination model and the tandem duplication and random loss (TDRL) model, and the tandem duplication and non-random loss (TDNL) model. In *S. latus,* the model of double replications and random loss has been proposed to account for the rearrangements (Shi et al., 2014). The genomic rearrangement could have functional implications, potentially affecting gene expression or protein interactions. For instance, gene rearrangement was found to be correlated with deep-sea adaptation in mussels (Zhang et al., 2021), corals (Wei et al., 2024) and caridean species (Kong et al., 2024). These make sense since mitochondria and mitogenomes play a pivotal role in aerobic respiration to generate energy, and deep sea has unique environmental characteristics including oxygen depletion and limited food availability. Further studies are needed to elucidate the functional consequences of the rearrangement in mitogenome of flatfish, especially for Samaridae family. In addition, some variations were observed in the non-protein coding regions, such as the dark red co-linear block, indicate a higher degree of evolutionary flexibility in these regions, which may be less constrained by functional constraints, allowing for a more rapid accumulation of mutations and structural changes.

One of the most central and fundamental objectives in biology is the reconstruction of the tree of life (Delsuc et al., 2005). Mitogenomes have been extensively used to infer the tempo and mode of evolutionary changes, as well as the processes underlying speciation and adaptive radiation. The phylogenetic relationships within and among flatfish families are still debated, with different studies proposing conflicting hypotheses (Betancur-R et al., 2014; Campbell et al., 2014; Duarte-Ribeiro et al., 2024; Lu et al., 2021). In this study, as stated in Methods and Results, the phylogenetic tree was constructed using two different methods, one is based on the complete mitogenomes and the other is only based on the concatenated PCGs. Both methods revealed a well-resolved evolutionary history, with major clades corresponding to taxonomic families and genera. Notably, using the whole genome information including gene arrangement, *S. latus* and *S. cristatus* form distinct outgroups, highlighting their uniqueness within the order. However, using concatenated independently aligned PCGs did not capture this information. In fact, many studies have emphasized the valuable information conveyed by gene order (Delsuc et al., 2005; Steenwyk and King, 2024), highlighting the importance of incorporating gene-order data in inferring phylogenies. It is worth noting that doubt should be cast on the use of gene order for phylogenetic analysis, since convergent evolution of mitochondrial gene order may occur. While our results point to the importance of whole mitogenome data, more extensive investigations are necessary before definitively concluding which method is more accurate.

The analysis of SSRs and various types of repeat sequences in the flatfish mitogenomes provides valuable insights into the genetic architecture and functional implication of these flatfish. SSRs play roles in DNA replication, transcription, mRNA splicing, translation, gene function and evolution (Gemayel et al., 2010; Haasl et al., 2012; Kashi and King, 2006; Li et al., 2004). Our comprehensive survey identified a total of 227 SSR sites, with monomeric repeats being the most abundant, followed by trimeric and tetrameric repeats. This distribution pattern suggests a preference for shorter, monomeric repeats within the mitochondrial DNA of flatfish, consistent with the findings in plants (Kuntal and Sharma, 2011). Based on the statistics in FMiR database (Nagpure et al., 2015), however, trinucleotide SSR is more dominant than any other SSR types in fish species. The observation emphasizes the unique characteristics of flatfish mitogenomes. The preponderance of T repeat monomers (59.6%) and the absence of (A)n further underscores the bias towards specific nucleotide repetitions in these mitogenomes. In addition to SSRs, the identification of forward, palindromic, reverse and complementary repeats add another layer of complexity to the genetic landscape of flatfish mitogenomes. They may affect replication, recombination and certain functions (Kurtz et al., 2001; Schon et al., 1989; Wynn and Christensen, 2019). For example, a 13-base pair repeat is responsible for large-scale deletion in the human mitogenome, leading to neuromuscular disorders including Kearns-Sayre syndrome and progressive external ophthalmoplegia (Schon et al., 1989). However, their specific functions in mitogenome remain largely underexplored in contrast to SSRs, and this disparity in understanding highlights a pivotal avenue for future research endeavors. Moreover, the abundance and diversity of SSRs and dispersed repeats in flatfish mitogenomes underscore their potential utility as molecular markers, aiding in conservation efforts, stock identification and selective breeding practices. SSRs, in particular, have been extensively used in genetic mapping, population genetics and breeding programs due to their high polymorphism and ease of genotyping (Vieira et al., 2016). Further investigation into the functional roles and population-level variation of these repeats will enhance our understanding of flatfish biology and contribute to the development of effective genetic management strategies.

While our study provides a comprehensive analysis of mitogenomes across 111 flatfish species, several limitations must be acknowledged. First, the use of complete mitogenomes, while powerful, may not capture the full complexity of evolutionary processes occurring at the nuclear level. Studies incorporating both mitochondrial and nuclear markers could provide a more holistic view of flatfish evolution. Second, further investigation should be performed for the species exhibiting unusual gene rearrangements to unravel the mechanisms driving these changes and their potential significance. Third, the functional roles of identified repeats remain to be experimentally examined. Future research is warranted to address these questions, paving the way for more comprehensive understanding and utilization.

## 5. Conclusions

In this study, using the mitochondrial genomes of 111 flatfish species, we studied the genomic structure, codon preference, nucleotide diversity, selective pressure and repeat sequences, as well as the phylogenetic relationship. This study revealed the relatively large gene rearrangement in Samaridae and Bothidae family, suggesting their unique evolutionary history and/or functional implications. Nucleotide diversity and selective pressure analysis indicated the conservation of *cytb* and *cox* genes, as well as the conservation levels of specific regions within genes. Phylogenetic analysis using different approaches exhibited different results, highlighting the role of incorporating gene order information in tree reconstruction. Furthermore, the identified repeats provided additional complexity of genome organization and offered valuable markers for evolutionary and functional studies. Our findings represent a significant step towards understanding the genetic architecture, evolutionary dynamics and functional implication of flatfish mitogenomes. The insights gained from this analysis have important implications for the study of adaptive evolution and fisheries management in marine organisms. With the continued advancements in multi-omics technologies, bioinformatics and validation methods, we anticipate that future research will further unravel the evolutionary history, functional implication and underlying mechanisms in these fascinating fish.

## CRediT authorship contribution statement

**Suxu Tan**: Conceptualization, Funding acquisition, Methodology, Data analysis, Writing - original draft, review and editing. **Wenwen Wang**: Writing - review and editing. **Jinjiang Li**: Methodology. **Zhenxia Sha**: Funding acquisition, Writing - review and editing.

## Availability of data and materials

All data will be made available on request.

## Acknowledgement

This study was supported by the National Key R&D Program of China (Grant No. 2022YFD2400401), Natural Science Foundation of Shandong Province (Grant No. ZR2023QC259), Shandong Key R&D Program for Academician team in Shandong (Grant No. 2023ZLYS02), and Taishan Scholar Youth Project of Shandong Province, China.

