## Supplementary material for "Comprehensive analysis of 111 Pleuronectiformes mitochondrial genomes: insights into structure, conservation, variation and evolution": Fig. S3

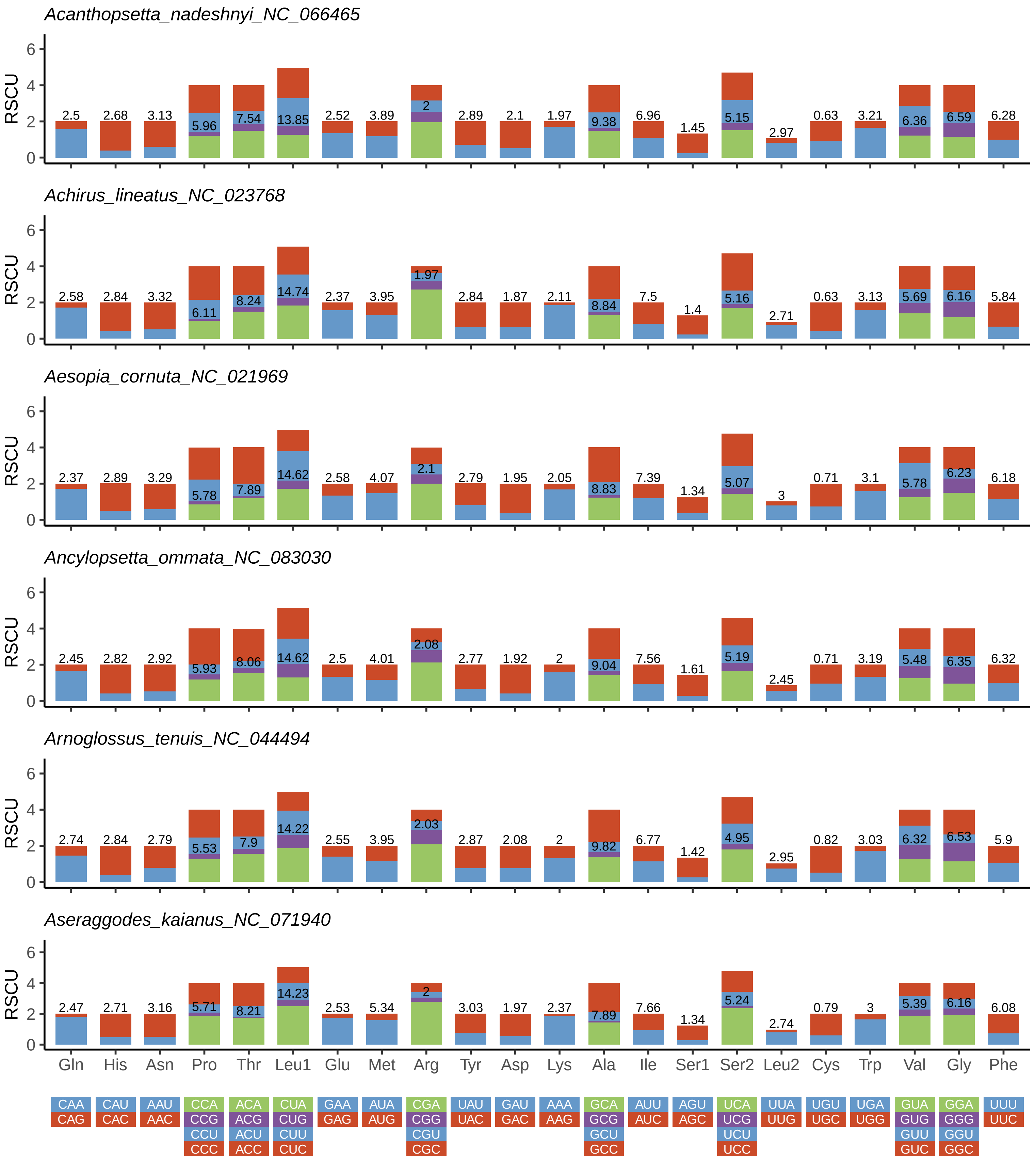
Acanthopsetta_nadeshnyi_NC_066465
2.5 2.68 3.13 5.96 7.54 13.85 2.52 3.89 2 2.89 2.1 1.97 9.38 6.96 1.45 5.15 2.97 0.63 3.21 6.36 6.59 6.28
Achirus_lineatus_NC_023768
2.58 2.84 3.32 6.11 8.24 14.74 2.37 3.95 1.97 2.84 1.87 2.11 8.84 7.5 1.4 5.16 2.71 0.63 3.13 5.69 6.16 5.84
Aesopia_cornuta_NC_021969
2.37 2.89 3.29 5.78 7.89 14.62 2.58 4.07 2.1 2.79 1.95 2.05 8.83 7.39 1.34 5.07 3 0.71 3.1 5.78 6.23 6.18
Ancylopsetta_ommata_NC_083030
2.45 2.82 2.92 5.93 8.06 14.62 2.5 4.01 2.08 2.77 1.92 2 9.04 7.56 1.61 5.19 2.45 0.71 3.19 5.48 6.35 6.32
Arnoglossus_tenuis_NC_044494
2.74 2.84 2.79 5.53 7.9 14.22 2.55 3.95 2.03 2.87 2.08 2 9.82 6.77 1.42 4.95 2.95 0.82 3.03 6.32 6.53 5.9
Aseraggodes_kaianus_NC_071940
2.47 2.71 3.16 5.71 8.21 14.23 2.53 5.34 2 3.03 1.97 2.37 7.89 7.66 1.34 5.24 2.74 0.79 3 5.39 6.16 6.08
RSCU
0 2 4 6
Gln His Asn Pro Thr Leu1 Glu Met Arg Tyr Asp Lys Ala Ile Ser1 Ser2 Leu2 Cys Trp Val Gly Phe
CAA CAU AAU CCA ACA CUA GAA AUA CGA UAU GAU AAA GCA AUU AGU UCA UUA UGU UGA GUA GGA UUU
CAG CAC AAC CCG ACG CUG GAG AUG CGG UAC GAC AAG GCG AUC AGC UCG UUG UGC UGG GUG GGG UUC
CCU ACU CUU CGU GCU UCU GUU GGU
CCC ACC CUC CGC GCC UCC GUC GGC

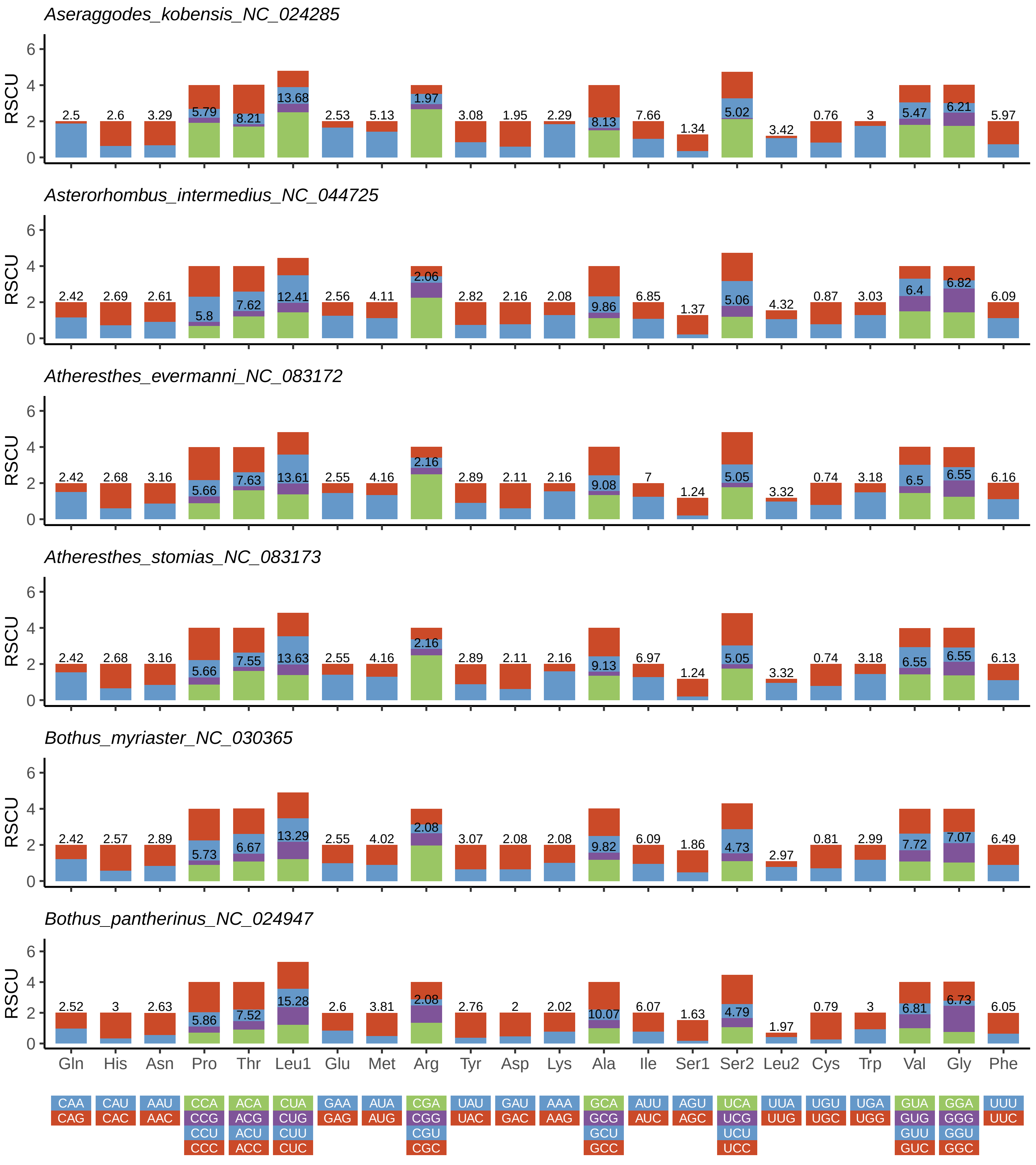

Aseraggodes_kobensis_NC_024285
Asterorhombus_intermedius_NC_044725
Atheresthes_evermanni_NC_083172
Atheresthes_stomias_NC_083173
Bothus_myriaster_NC_030365
Bothus_pantherinus_NC_024947
RSCU
Gln His Asn Pro Thr Leu1 Glu Met Arg Tyr Asp Lys Ala Ile Ser1 Ser2 Leu2 Cys Trp Val Gly Phe
CAA CAG
CAU CAC
AAU AAC
CCA CCG CCU CCC
ACA ACG ACU ACC
CUA CUG CUU CUC
GAA GAG
AUA AUG
CGA CGG CGU CGC
UAU UAC
GAU GAC
AAA AAG
GCA GCG GCU GCC
AUU AUC
AGU AGC
UCA UCG UCU UCC
UUA UUG
UGU UGC
UGA UGG
GUA GUG GUU GUC
GGA GGG GGU GGC
UUU UUC

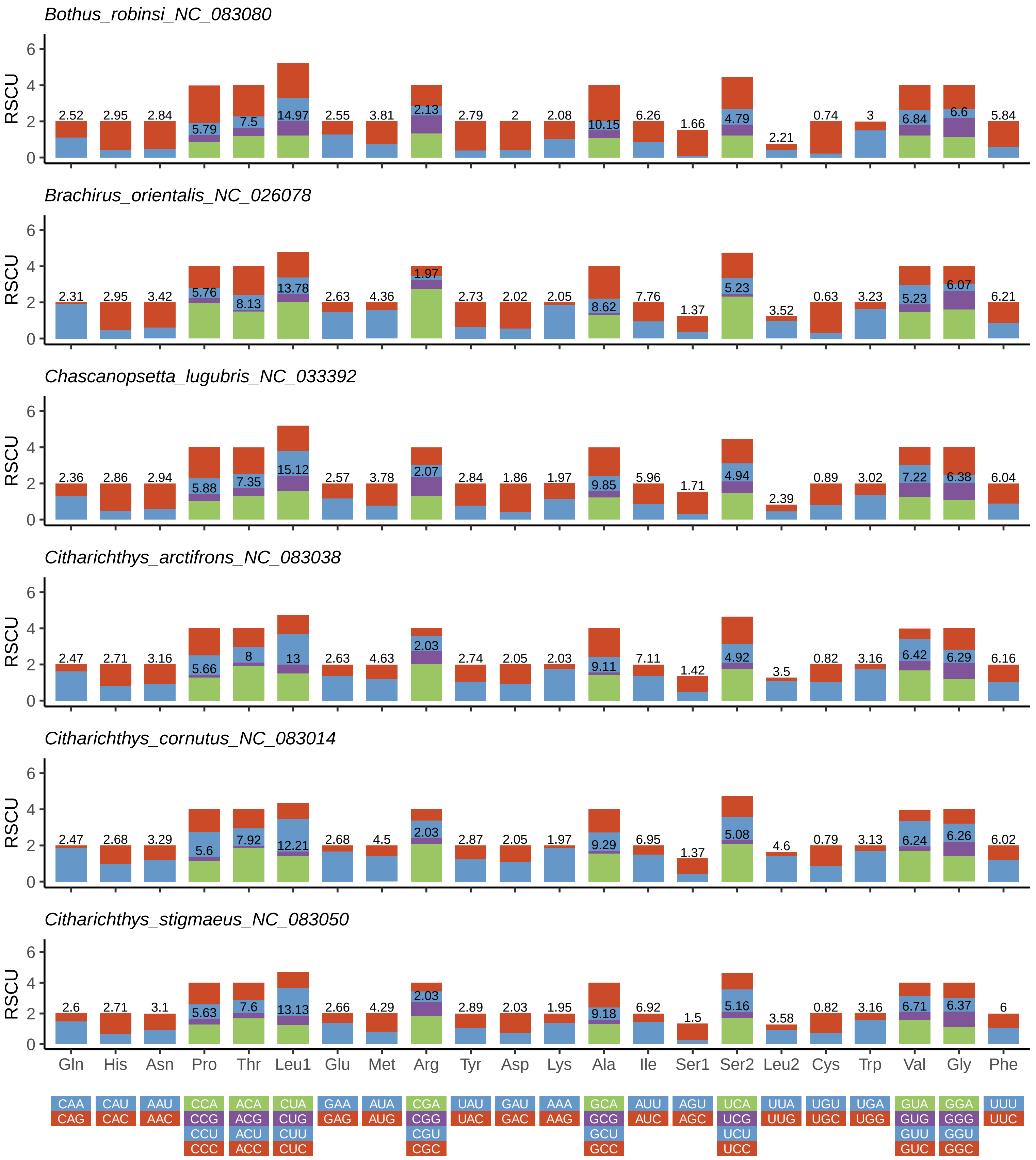
Bothus_robinsi_NC_083080
RSCU
0 2 4 6
2.52 2.95 2.84 5.79 7.5 14.97 2.55 3.81 2.13 2.79 2 2.08 10.15 6.26 1.66 4.79 2.21 0.74 3 6.84 6.6 5.84
Brachirus_orientalis_NC_026078
RSCU
0 2 4 6
2.31 2.95 3.42 5.76 8.13 13.78 2.63 4.36 1.97 2.73 2.02 2.05 8.62 7.76 1.37 5.23 3.52 0.63 3.23 5.23 6.07 6.21
Chascanopsetta_lugubris_NC_033392
RSCU
0 2 4 6
2.36 2.86 2.94 5.88 7.35 15.12 2.57 3.78 2.07 2.84 1.86 1.97 9.85 5.96 1.71 4.94 2.39 0.89 3.02 7.22 6.38 6.04
Citharichthys_arctifrons_NC_083038
RSCU
0 2 4 6
2.47 2.71 3.16 5.66 8 13 2.63 4.63 2.03 2.74 2.05 2.03 9.11 7.11 1.42 4.92 3.5 0.82 3.16 6.42 6.29 6.16
Citharichthys_cornutus_NC_083014
RSCU
0 2 4 6
2.47 2.68 3.29 5.6 7.92 12.21 2.68 4.5 2.03 2.87 2.05 1.97 9.29 6.95 1.37 5.08 4.6 0.79 3.13 6.24 6.26 6.02
Citharichthys_stigmaeus_NC_083050
RSCU
0 2 4 6
2.6 2.71 3.1 5.63 7.6 13.13 2.66 4.29 2.03 2.89 2.03 1.95 9.18 6.92 1.5 5.16 3.58 0.82 3.16 6.71 6.37 6
Gln His Asn Pro Thr Leu1 Glu Met Arg Tyr Asp Lys Ala Ile Ser1 Ser2 Leu2 Cys Trp Val Gly Phe
CAA CAU AAU CCA ACA CUA GAA AUA CGA UAU GAU AAA GCA AUU AGU UCA UUA UGU UGA GUA GGA UUU
CAG CAC AAC CCG ACG CUG GAG AUG CGG UAC GAC AAG GCG AUC AGC UCG UUG UGC UGG GUG GGG UUC
CCU ACU CUU CGU GCU UCU GUU GGU
CCC ACC CUC CGC GCC UCC GUC GGC

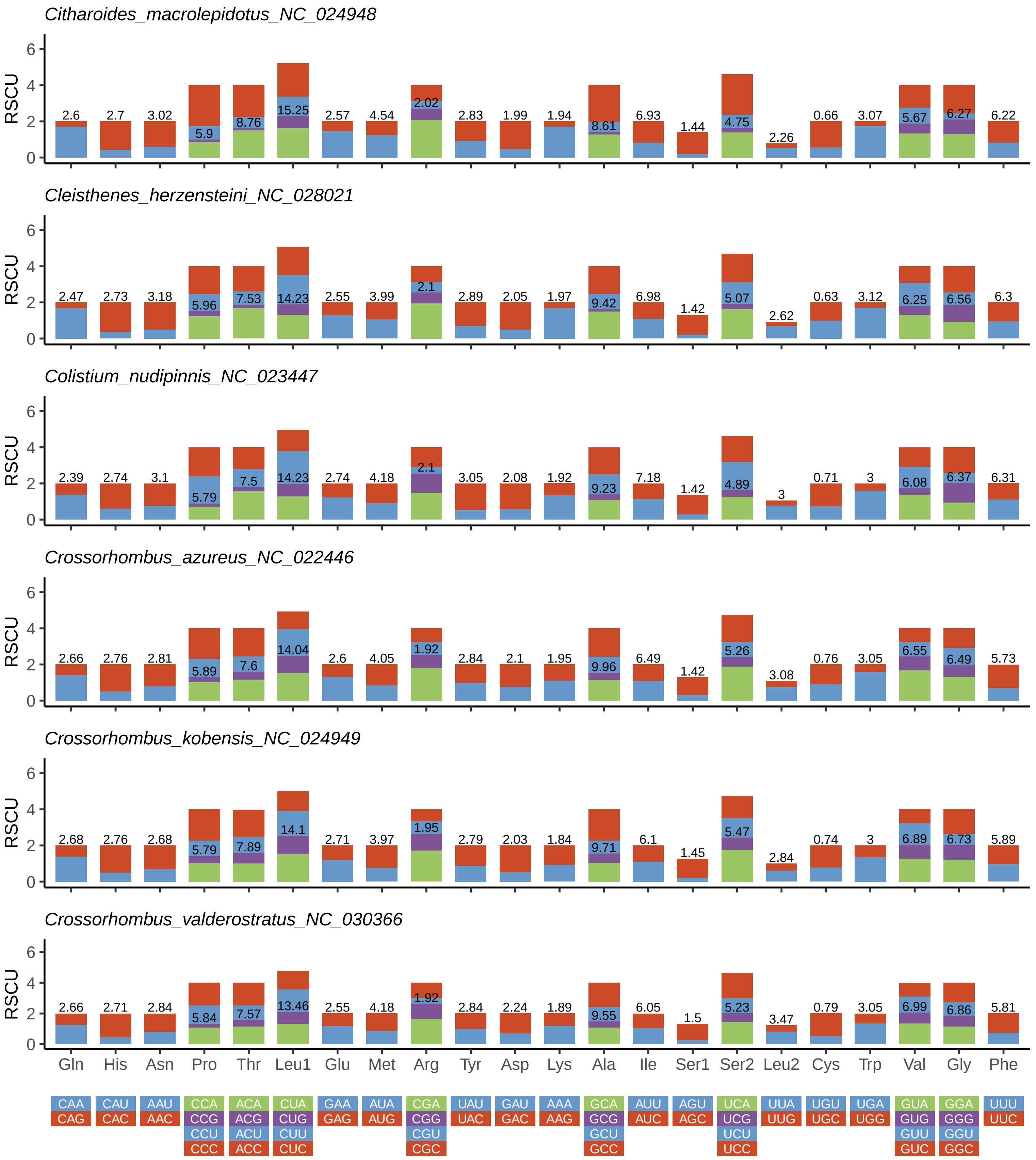

Citharoides_macrolepidotus_NC_024948
RSCU
0 2 4 6
2.6 2.7 3.02 5.9 8.76 15.25 2.57 4.54 2.02 2.83 1.99 1.94 8.61 6.93 1.44 4.75 2.26 0.66 3.07 5.67 6.27 6.22
Cleisthenes_herzensteini_NC_028021
RSCU
0 2 4 6
2.47 2.73 3.18 5.96 7.53 14.23 2.55 3.99 2.1 2.89 2.05 1.97 9.42 6.98 1.42 5.07 2.62 0.63 3.12 6.25 6.56 6.3
Colistium_nudipinnis_NC_023447
RSCU
0 2 4 6
2.39 2.74 3.1 5.79 7.5 14.23 2.74 4.18 2.1 3.05 2.08 1.92 9.23 7.18 1.42 4.89 3 0.71 3 6.08 6.37 6.31
Crossorhombus_azureus_NC_022446
RSCU
0 2 4 6
2.66 2.76 2.81 5.89 7.6 14.04 2.6 4.05 1.92 2.84 2.1 1.95 9.96 6.49 1.42 5.26 3.08 0.76 3.05 6.55 6.49 5.73
Crossorhombus_kobensis_NC_024949
RSCU
0 2 4 6
2.68 2.76 2.68 5.79 7.89 14.1 2.71 3.97 1.95 2.79 2.03 1.84 9.71 6.1 1.45 5.47 2.84 0.74 3 6.89 6.73 5.89
Crossorhombus_valderostratus_NC_030366
RSCU
0 2 4 6
2.66 2.71 2.84 5.84 7.57 13.46 2.55 4.18 1.92 2.84 2.24 1.89 9.55 6.05 1.5 5.23 3.47 0.79 3.05 6.99 6.86 5.81
Gln His Asn Pro Thr Leu1 Glu Met Arg Tyr Asp Lys Ala Ile Ser1 Ser2 Leu2 Cys Trp Val Gly Phe
CAA CAU AAU CCA ACA CUA GAA AUA CGA UAU GAU AAA GCA AUU AGU UCA UUA UGU UGA GUA GGA UUU
CAG CAC AAC CCG ACG CUG GAG AUG CGG UAC GAC AAG GCG AUC AGC UCG UUG UGC UGG GUG GGG UUC
CCU ACU CUU CGU GCU UCU GUU GGU
CCC ACC CUC CGC GCC UCC GUC GGC

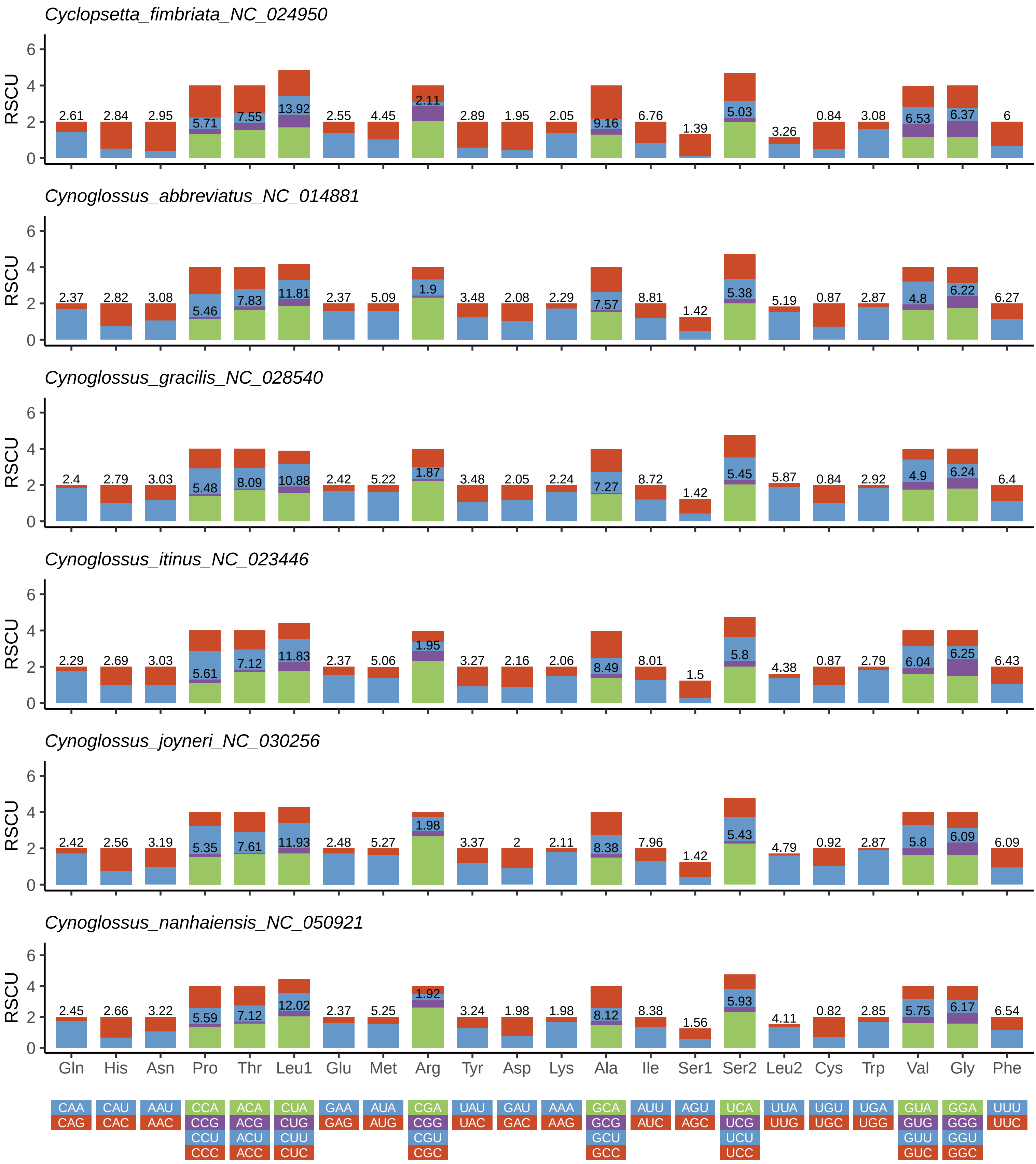
Cyclopsetta_fimbriata_NC_024950
RSCU
2.61 2.84 2.95 5.71 7.55 13.92 2.55 4.45 2.11 2.89 1.95 2.05 9.16 6.76 1.39 5.03 3.26 0.84 3.08 6.53 6.37 6
Cynoglossus_abbreviatus_NC_014881
RSCU
2.37 2.82 3.08 5.46 7.83 11.81 2.37 5.09 1.9 3.48 2.08 2.29 7.57 8.81 1.42 5.38 5.19 0.87 2.87 4.8 6.22 6.27
Cynoglossus_gracilis_NC_028540
RSCU
2.4 2.79 3.03 5.48 8.09 10.88 2.42 5.22 1.87 3.48 2.05 2.24 7.27 8.72 1.42 5.45 5.87 0.84 2.92 4.9 6.24 6.4
Cynoglossus_itinus_NC_023446
RSCU
2.29 2.69 3.03 5.61 7.12 11.83 2.37 5.06 1.95 3.27 2.16 2.06 8.49 8.01 1.5 5.8 4.38 0.87 2.79 6.04 6.25 6.43
Cynoglossus_joyneri_NC_030256
RSCU
2.42 2.56 3.19 5.35 7.61 11.93 2.48 5.27 1.98 3.37 2 2.11 8.38 7.96 1.42 5.43 4.79 0.92 2.87 5.8 6.09 6.09
Cynoglossus_nanhaiensis_NC_050921
RSCU
2.45 2.66 3.22 5.59 7.12 12.02 2.37 5.25 1.92 3.24 1.98 1.98 8.12 8.38 1.56 5.93 4.11 0.82 2.85 5.75 6.17 6.54
0 2 4 6
Gln His Asn Pro Thr Leu1 Glu Met Arg Tyr Asp Lys Ala Ile Ser1 Ser2 Leu2 Cys Trp Val Gly Phe
CAA CAU AAU CCA ACA CUA GAA AUA CGA UAU GAU AAA GCA AUU AGU UCA UUA UGU UGA GUA GGA UUU
CAG CAC AAC CCG ACG CUG GAG AUG CGG UAC GAC AAG GCG AUC AGC UCG UUG UGC UGG GUG GGG UUC
CCU ACU CUU CGU GCU UCU GUU GGU
CCC ACC CUC CGC GCC UCC GUC GGC

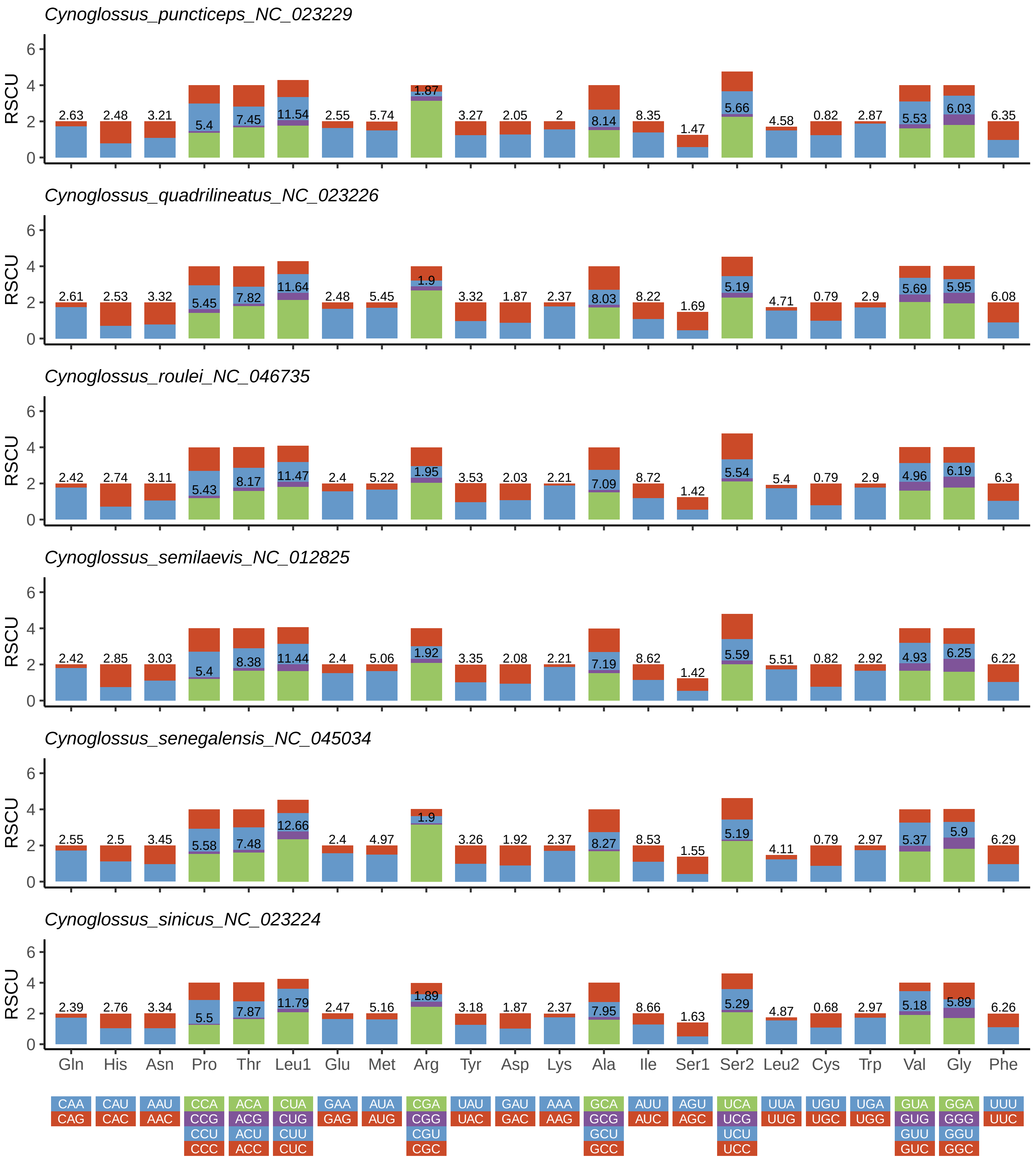
Cynoglossus_puncticeps_NC_023229
Cynoglossus_quadrilineatus_NC_023226
Cynoglossus_roulei_NC_046735
Cynoglossus_semilaevis_NC_012825
Cynoglossus_senegalensis_NC_045034
Cynoglossus_sinicus_NC_023224
RSCU
Gln His Asn Pro Thr Leu1 Glu Met Arg Tyr Asp Lys Ala Ile Ser1 Ser2 Leu2 Cys Trp Val Gly Phe
CAA CAU AAU CCA ACA CUA GAA AUA CGA UAU GAU AAA GCA AUU AGU UCA UUA UGU UGA GUA GGA UUU
CAG CAC AAC CCG ACG CUG GAG AUG CGG UAC GAC AAG GCG AUC AGC UCG UUG UGC UGG GUG GGG UUC
CCU ACU CUU CGU GCU UCU GUU GGU
CCC ACC CUC CGC GCC UCC GUC GGC

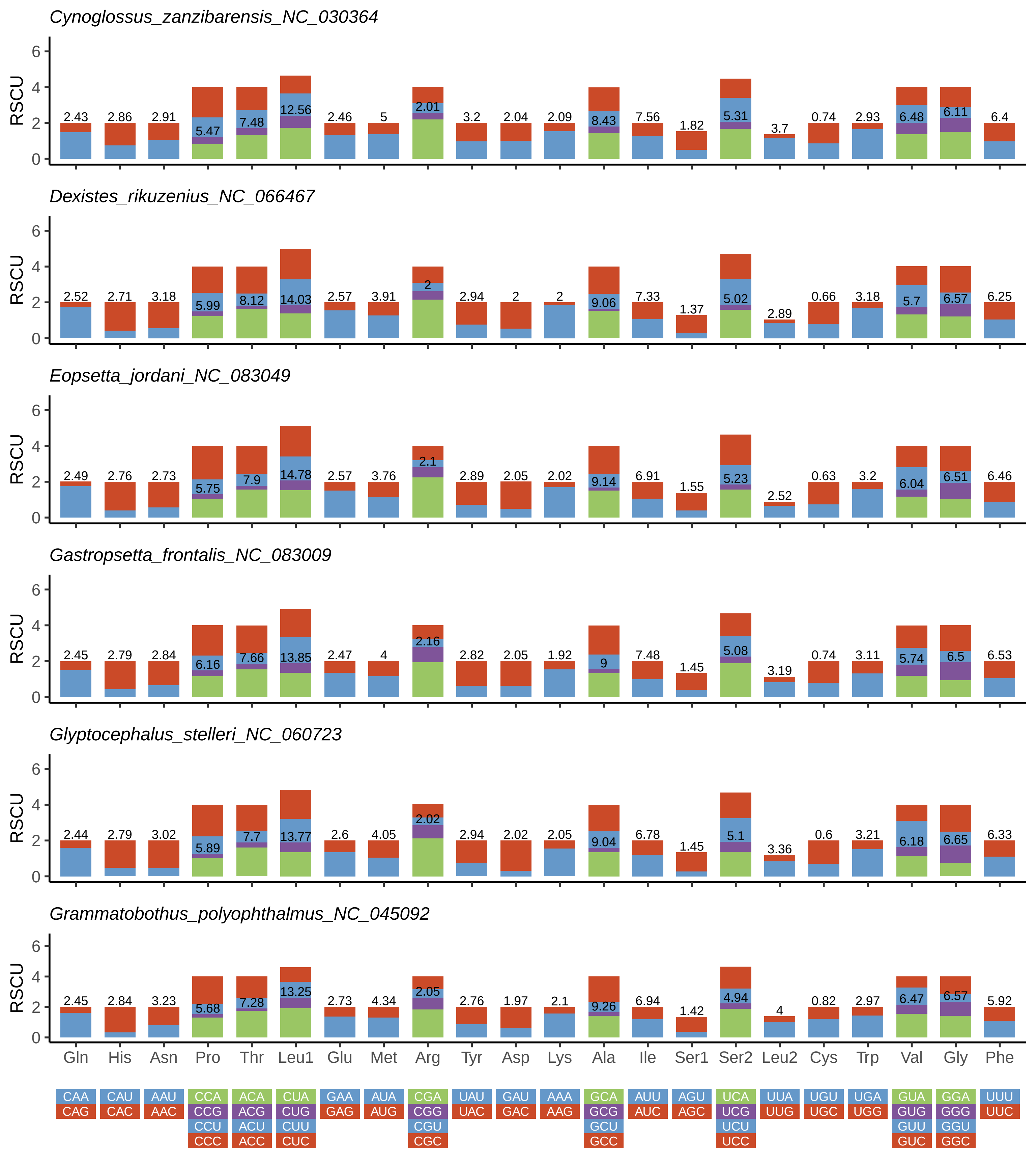

Cynoglossus_zanzibarensis_NC_030364
RSCU
2.43 2.86 2.91 5.47 7.48 12.56 2.46 5 2.01 3.2 2.04 2.09 8.43 7.56 1.82 5.31 3.7 0.74 2.93 6.48 6.11 6.4
Dexistes_rikuzenius_NC_066467
RSCU
2.52 2.71 3.18 5.99 8.12 14.03 2.57 3.91 2 2.94 2 2 9.06 7.33 1.37 5.02 2.89 0.66 3.18 5.7 6.57 6.25
Eopsetta_jordani_NC_083049
RSCU
2.49 2.76 2.73 5.75 7.9 14.78 2.57 3.76 2.1 2.89 2.05 2.02 9.14 6.91 1.55 5.23 2.52 0.63 3.2 6.04 6.51 6.46
Gastropsetta_frontalis_NC_083009
RSCU
2.45 2.79 2.84 6.16 7.66 13.85 2.47 4 2.16 2.82 2.05 1.92 9 7.48 1.45 5.08 3.19 0.74 3.11 5.74 6.5 6.53
Glyptocephalus_stelleri_NC_060723
RSCU
2.44 2.79 3.02 5.89 7.7 13.77 2.6 4.05 2.02 2.94 2.02 2.05 9.04 6.78 1.45 5.1 3.36 0.6 3.21 6.18 6.65 6.33
Grammatobothus_polyophthalmus_NC_045092
RSCU
2.45 2.84 3.23 5.68 7.28 13.25 2.73 4.34 2.05 2.76 1.97 2.1 9.26 6.94 1.42 4.94 4 0.82 2.97 6.47 6.57 5.92
0 2 4 6
Gln His Asn Pro Thr Leu1 Glu Met Arg Tyr Asp Lys Ala Ile Ser1 Ser2 Leu2 Cys Trp Val Gly Phe
CAA CAU AAU CCA ACA CUA GAA AUA CGA UAU GAU AAA GCA AUU AGU UCA UUA UGU UGA GUA GGA UUU
CAG CAC AAC CCG ACG CUG GAG AUG CGG UAC GAC AAG GCG AUC AGC UCG UUG UGC UGG GUG GGG UUC
CCU ACU CUU CGU GCU UCU GUU GGU
CCC ACC CUC CGC GCC UCC GUC GGC

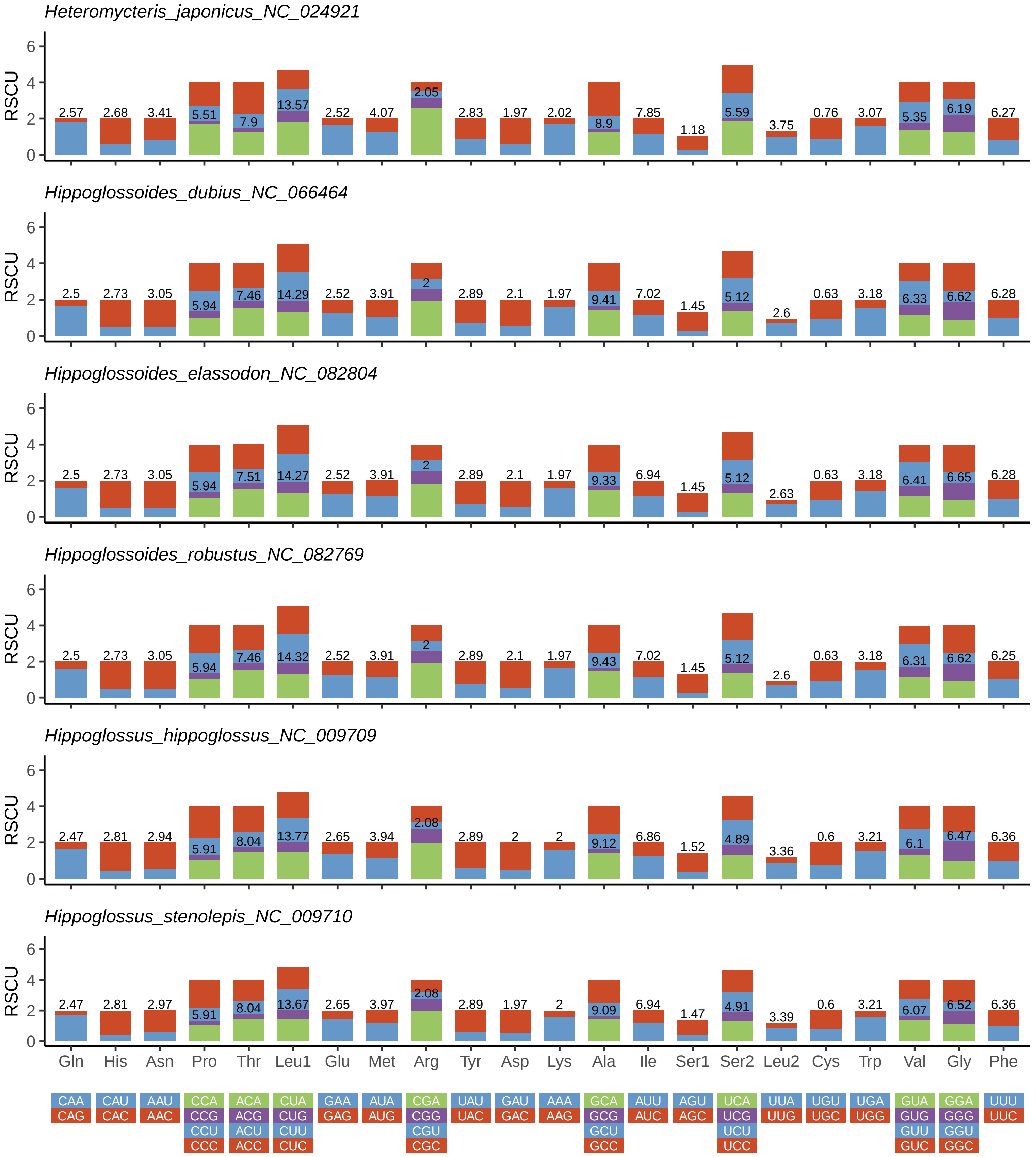

Heteromycteris_japonicus_NC_024921
Hippoglossoides_dubius_NC_066464
Hippoglossoides_elassodon_NC_082804
Hippoglossoides_robustus_NC_082769
Hippoglossus_hippoglossus_NC_009709
Hippoglossus_stenolepis_NC_009710
RSCU
Gln
His
Asn
Pro
Thr
Leu1
Glu
Met
Arg
Tyr
Asp
Lys
Ala
Ile
Ser1
Ser2
Leu2
Cys
Trp
Val
Gly
Phe
CAA CAG
CAU CAC
AAU AAC
CCA CCG CCU CCC
ACA ACG ACU ACC
CUA CUG CUU CUC
GAA GAG
AUA AUG
CGA CGG CGU CGC
UAU UAC
GAU GAC
AAA AAG
GCA GCG GCU GCC
AUU AUC
AGU AGC
UCA UCG UCU UCC
UUA UUG
UGU UGC
UGA UGG
GUA GUG GUU GUC
GGA GGG GGU GGC
UUU UUC

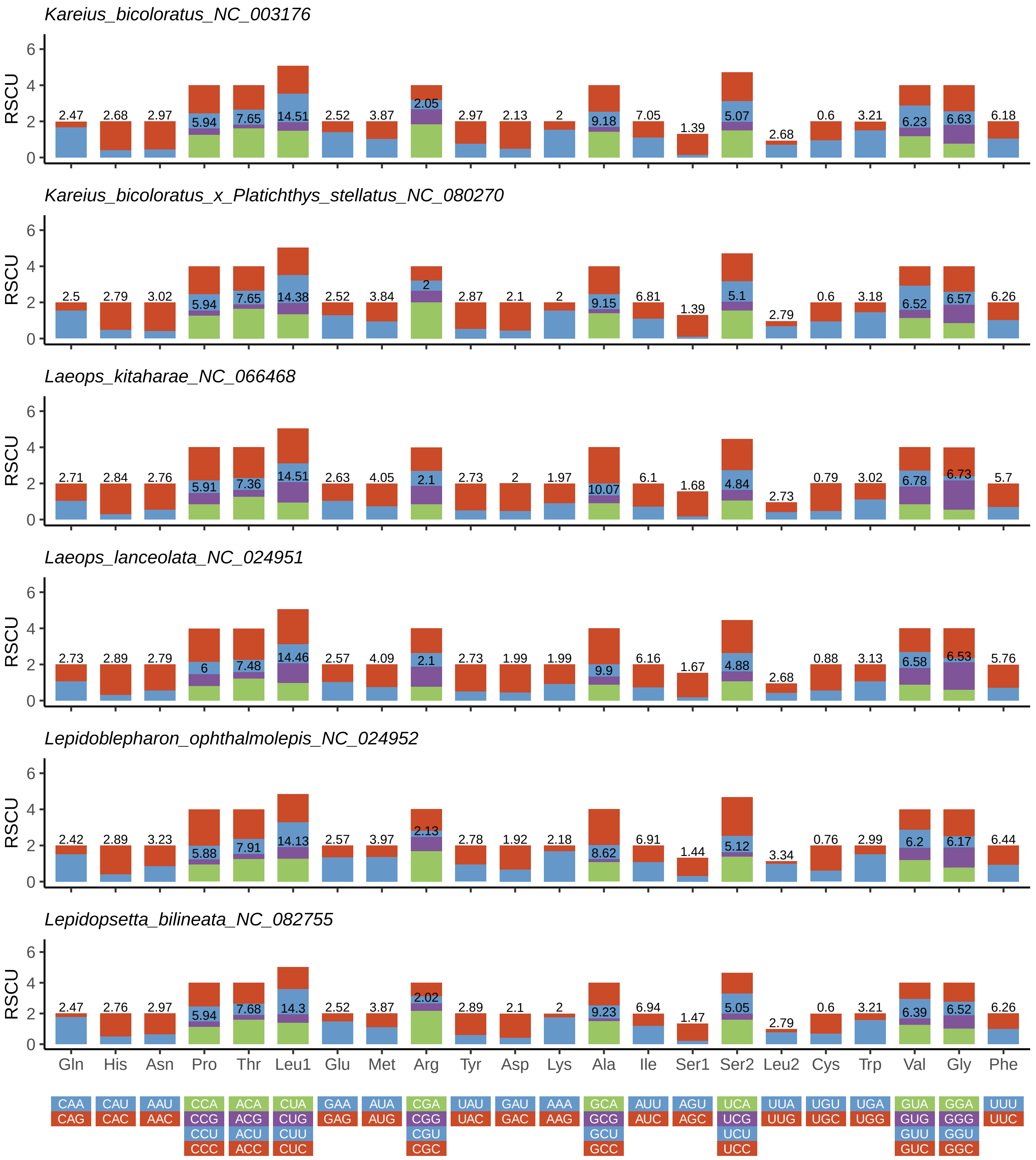

Kareius_bicoloratus_NC_003176
Kareius_bicoloratus_x_Platichthys_stellatus_NC_080270
Laeops_kitaharae_NC_066468
Laeops_lanceolata_NC_024951
Lepidoblepharon_ophthalmolepis_NC_024952
Lepidopsetta_bilineata_NC_082755
RSCU
Gln His Asn Pro Thr Leu1 Glu Met Arg Tyr Asp Lys Ala Ile Ser1 Ser2 Leu2 Cys Trp Val Gly Phe
CAA CAG
CAU CAC
AAU AAC
CCA CCG CCU CCC
ACA ACG ACU ACC
CUA CUG CUU CUC
GAA GAG
AUA AUG
CGA CGG CGU CGC
UAU UAC
GAU GAC
AAA AAG
GCA GCG GCU GCC
AUU AUC
AGU AGC
UCA UCG UCU UCC
UUA UUG
UGU UGC
UGA UGG
GUA GUG GUU GUC
GGA GGG GGU GGC
UUU UUC

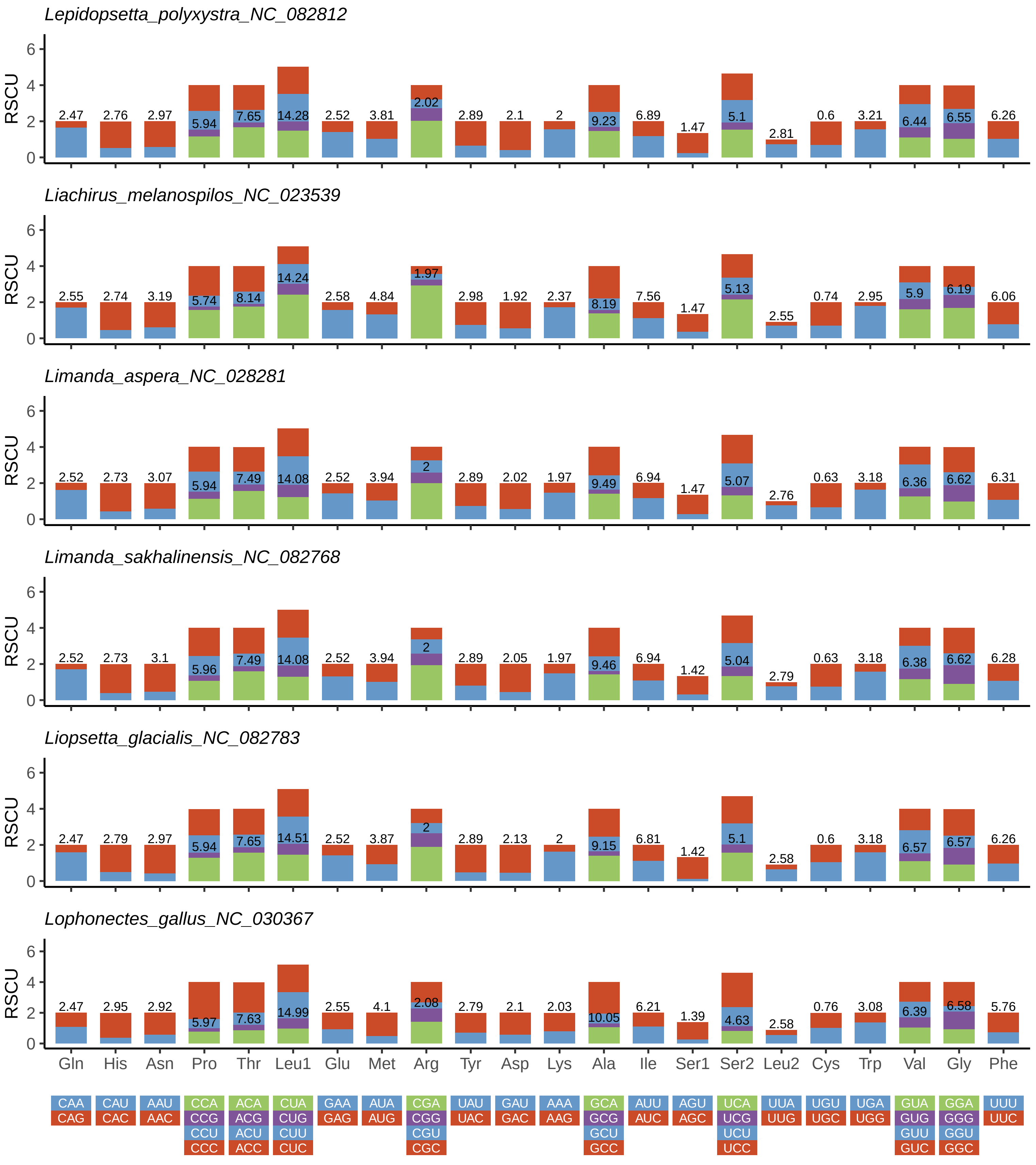
Lepidopsetta_polyxystra_NC_082812
RSCU
0 2 4 6
2.47 2.76 2.97 5.94 7.65 14.28 2.52 3.81 2.02 2.89 2.1 2 9.23 6.89 1.47 5.1 2.81 0.6 3.21 6.44 6.55 6.26
Liachirus_melanospilos_NC_023539
RSCU
0 2 4 6
2.55 2.74 3.19 5.74 8.14 14.24 2.58 4.84 1.97 2.98 1.92 2.37 8.19 7.56 1.47 5.13 2.55 0.74 2.95 5.9 6.19 6.06
Limanda_aspera_NC_028281
RSCU
0 2 4 6
2.52 2.73 3.07 5.94 7.49 14.08 2.52 3.94 2 2.89 2.02 1.97 9.49 6.94 1.47 5.07 2.76 0.63 3.18 6.36 6.62 6.31
Limanda_sakhalinensis_NC_082768
RSCU
0 2 4 6
2.52 2.73 3.1 5.96 7.49 14.08 2.52 3.94 2 2.89 2.05 1.97 9.46 6.94 1.42 5.04 2.79 0.63 3.18 6.38 6.62 6.28
Liopsetta_glacialis_NC_082783
RSCU
0 2 4 6
2.47 2.79 2.97 5.94 7.65 14.51 2.52 3.87 2 2.89 2.13 2 9.15 6.81 1.42 5.1 2.58 0.6 3.18 6.57 6.57 6.26
Lophonectes_gallus_NC_030367
RSCU
0 2 4 6
2.47 2.95 2.92 5.97 7.63 14.99 2.55 4.1 2.08 2.79 2.1 2.03 10.05 6.21 1.39 4.63 2.58 0.76 3.08 6.39 6.58 5.76
Gln His Asn Pro Thr Leu1 Glu Met Arg Tyr Asp Lys Ala Ile Ser1 Ser2 Leu2 Cys Trp Val Gly Phe
CAA CAG
CAU CAC
AAU AAC
CCA CCG CCU CCC
ACA ACG ACU ACC
CUA CUG CUU CUC
GAA GAG
AUA AUG
CGA CGG CGU CGC
UAU UAC
GAU GAC
AAA AAG
GCA GCG GCU GCC
AUU AUC
AGU AGC
UCA UCG UCU UCC
UUA UUG
UGU UGC
UGA UGG
GUA GUG GUU GUC
GGA GGG GGU GGC
UUU UUC

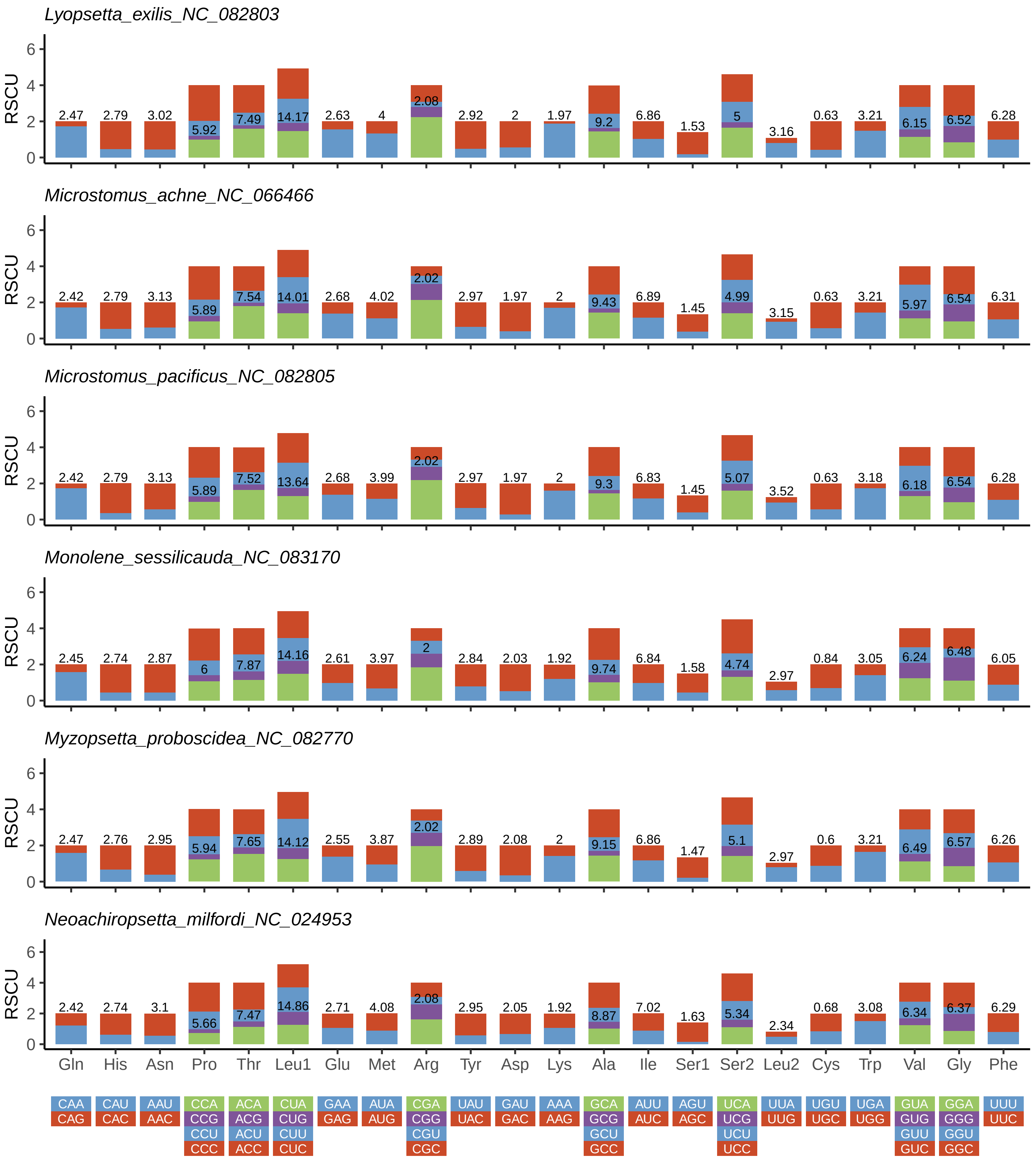

Lyopsetta_exilis_NC_082803
RSCU
0 2 4 6
2.47 2.79 3.02 5.92 7.49 14.17 2.63 4 2.08 2.92 2 1.97 9.2 6.86 1.53 5 3.16 0.63 3.21 6.15 6.52 6.28
Microstomus_achne_NC_066466
RSCU
0 2 4 6
2.42 2.79 3.13 5.89 7.54 14.01 2.68 4.02 2.02 2.97 1.97 2 9.43 6.89 1.45 4.99 3.15 0.63 3.21 5.97 6.54 6.31
Microstomus_pacificus_NC_082805
RSCU
0 2 4 6
2.42 2.79 3.13 5.89 7.52 13.64 2.68 3.99 2.02 2.97 1.97 2 9.3 6.83 1.45 5.07 3.52 0.63 3.18 6.18 6.54 6.28
Monolene_sessilicauda_NC_083170
RSCU
0 2 4 6
2.45 2.74 2.87 6 7.87 14.16 2.61 3.97 2 2.84 2.03 1.92 9.74 6.84 1.58 4.74 2.97 0.84 3.05 6.24 6.48 6.05
Myzopsetta_proboscidea_NC_082770
RSCU
0 2 4 6
2.47 2.76 2.95 5.94 7.65 14.12 2.55 3.87 2.02 2.89 2.08 2 9.15 6.86 1.47 5.1 2.97 0.6 3.21 6.49 6.57 6.26
Neoachiropsetta_milfordi_NC_024953
RSCU
0 2 4 6
2.42 2.74 3.1 5.66 7.47 14.86 2.71 4.08 2.08 2.95 2.05 1.92 8.87 7.02 1.63 5.34 2.34 0.68 3.08 6.34 6.37 6.29
Gln His Asn Pro Thr Leu1 Glu Met Arg Tyr Asp Lys Ala Ile Ser1 Ser2 Leu2 Cys Trp Val Gly Phe
CAA CAU AAU CCA ACA CUA GAA AUA CGA UAU GAU AAA GCA AUU AGU UCA UUA UGU UGA GUA GGA UUU
CAG CAC AAC CCG ACG CUG GAG AUG CGG UAC GAC AAG GCG AUC AGC UCG UUG UGC UGG GUG GGG UUC
CCU ACU CUU CGU GCU UCU GUU GGU
CCC ACC CUC CGC GCC UCC GUC GGC

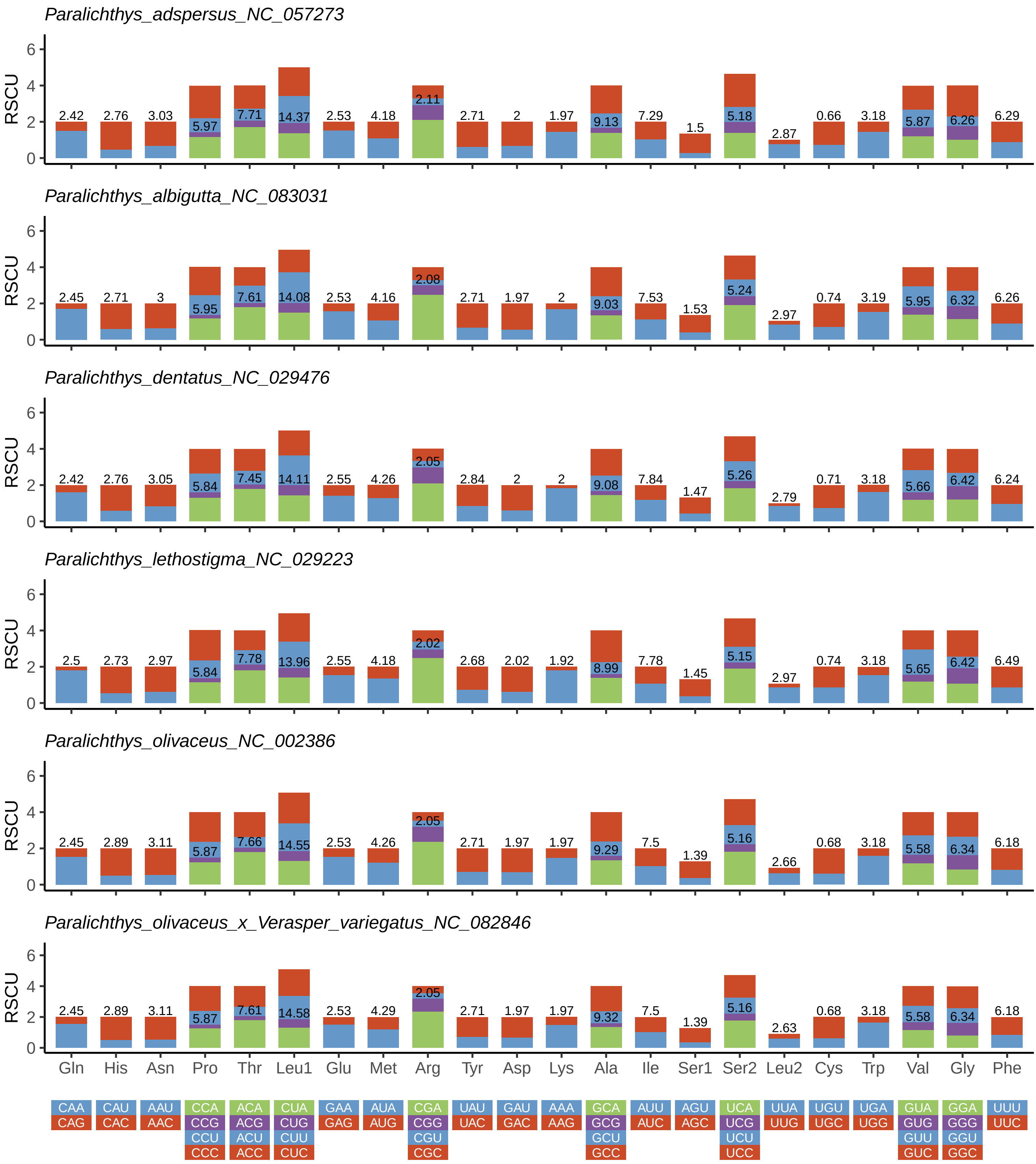

Paralichthys_adspersus_NC_057273
2.42 2.76 3.03 5.97 7.71 14.37 2.53 4.18 2.11 2.71 2 1.97 9.13 7.29 1.5 5.18 2.87 0.66 3.18 5.87 6.26 6.29
Paralichthys_albigutta_NC_083031
2.45 2.71 3 5.95 7.61 14.08 2.53 4.16 2.08 2.71 1.97 2 9.03 7.53 1.53 5.24 2.97 0.74 3.19 5.95 6.32 6.26
Paralichthys_dentatus_NC_029476
2.42 2.76 3.05 5.84 7.45 14.11 2.55 4.26 2.05 2.84 2 2 9.08 7.84 1.47 5.26 2.79 0.71 3.18 5.66 6.42 6.24
Paralichthys_lethostigma_NC_029223
2.5 2.73 2.97 5.84 7.78 13.96 2.55 4.18 2.02 2.68 2.02 1.92 8.99 7.78 1.45 5.15 2.97 0.74 3.18 5.65 6.42 6.49
Paralichthys_olivaceus_NC_002386
2.45 2.89 3.11 5.87 7.66 14.55 2.53 4.26 2.05 2.71 1.97 1.97 9.29 7.5 1.39 5.16 2.66 0.68 3.18 5.58 6.34 6.18
Paralichthys_olivaceus_x_Verasper_variegatus_NC_082846
2.45 2.89 3.11 5.87 7.61 14.58 2.53 4.29 2.05 2.71 1.97 1.97 9.32 7.5 1.39 5.16 2.63 0.68 3.18 5.58 6.34 6.18
RSCU
0 2 4 6
Gln His Asn Pro Thr Leu1 Glu Met Arg Tyr Asp Lys Ala Ile Ser1 Ser2 Leu2 Cys Trp Val Gly Phe
CAA CAU AAU CCA ACA CUA GAA AUA CGA UAU GAU AAA GCA AUU AGU UCA UUA UGU UGA GUA GGA UUU
CAG CAC AAC CCG ACG CUG GAG AUG CGG UAC GAC AAG GCG AUC AGC UCG UUG UGC UGG GUG GGG UUC
CCU ACU CUU CGU GCU UCU GUU GGU
CCC ACC CUC CGC GCC UCC GUC GGC

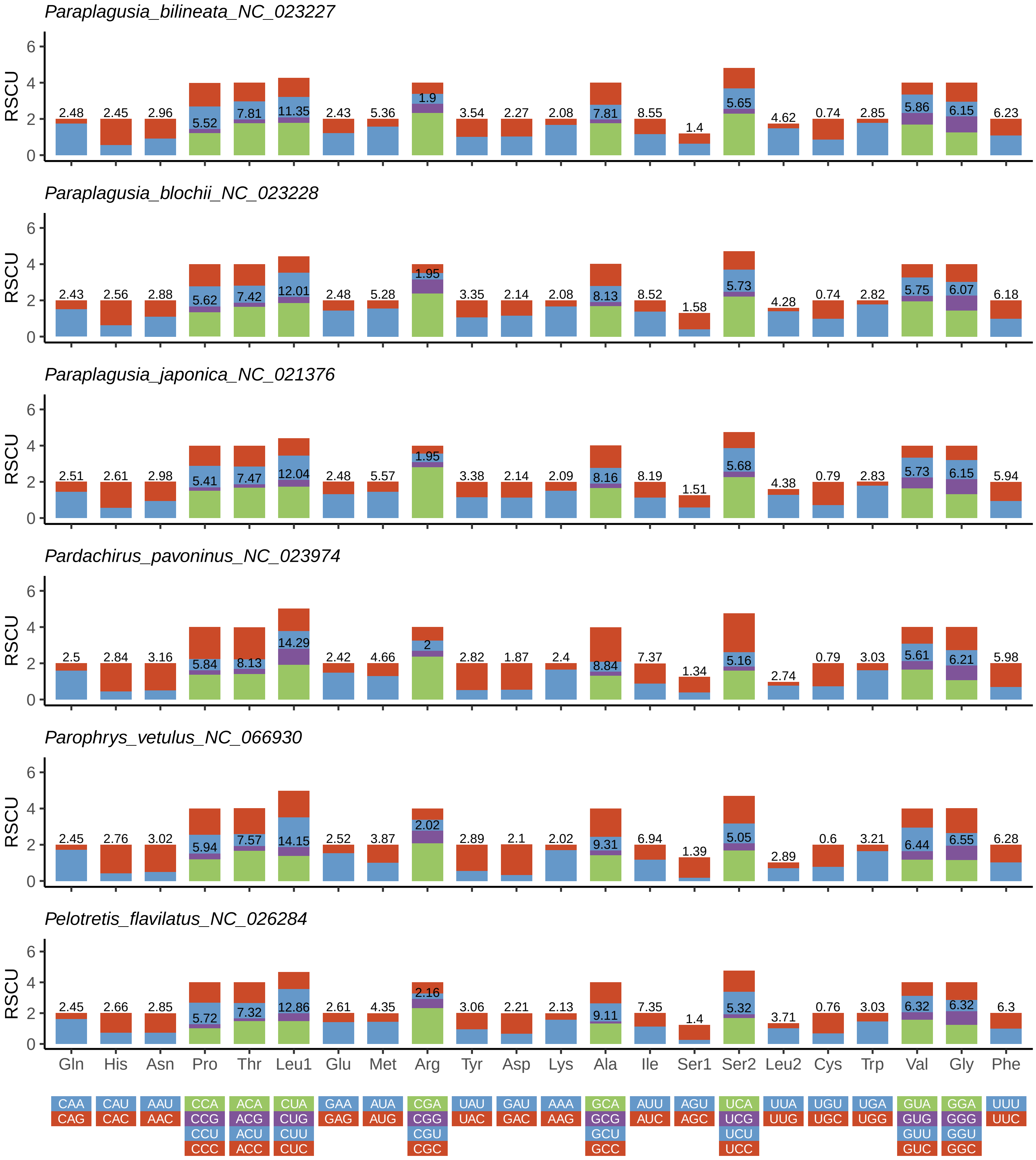
Paraplagusia_bilineata_NC_023227
RSCU
2.48 2.45 2.96 5.52 7.81 11.35 2.43 5.36 1.9 3.54 2.27 2.08 7.81 8.55 1.4 5.65 4.62 0.74 2.85 5.86 6.15 6.23
Paraplagusia_blochii_NC_023228
RSCU
2.43 2.56 2.88 5.62 7.42 12.01 2.48 5.28 1.95 3.35 2.14 2.08 8.13 8.52 1.58 5.73 4.28 0.74 2.82 5.75 6.07 6.18
Paraplagusia_japonica_NC_021376
RSCU
2.51 2.61 2.98 5.41 7.47 12.04 2.48 5.57 1.95 3.38 2.14 2.09 8.16 8.19 1.51 5.68 4.38 0.79 2.83 5.73 6.15 5.94
Pardachirus_pavoninus_NC_023974
RSCU
2.5 2.84 3.16 5.84 8.13 14.29 2.42 4.66 2 2.82 1.87 2.4 8.84 7.37 1.34 5.16 2.74 0.79 3.03 5.61 6.21 5.98
Parophrys_vetulus_NC_066930
RSCU
2.45 2.76 3.02 5.94 7.57 14.15 2.52 3.87 2.02 2.89 2.1 2.02 9.31 6.94 1.39 5.05 2.89 0.6 3.21 6.44 6.55 6.28
Pelotretis_flavilatus_NC_026284
RSCU
2.45 2.66 2.85 5.72 7.32 12.86 2.61 4.35 2.16 3.06 2.21 2.13 9.11 7.35 1.4 5.32 3.71 0.76 3.03 6.32 6.32 6.3
0 2 4 6
Gln His Asn Pro Thr Leu1 Glu Met Arg Tyr Asp Lys Ala Ile Ser1 Ser2 Leu2 Cys Trp Val Gly Phe
CAA CAU AAU CCA ACA CUA GAA AUA CGA UAU GAU AAA GCA AUU AGU UCA UUA UGU UGA GUA GGA UUU
CAG CAC AAC CCG ACG CUG GAG AUG CGG UAC GAC AAG GCG AUC AGC UCG UUG UGC UGG GUG GGG UUC
CCU ACU CUU CGU GCU UCU GUU GGU
CCC ACC CUC CGC GCC UCC GUC GGC

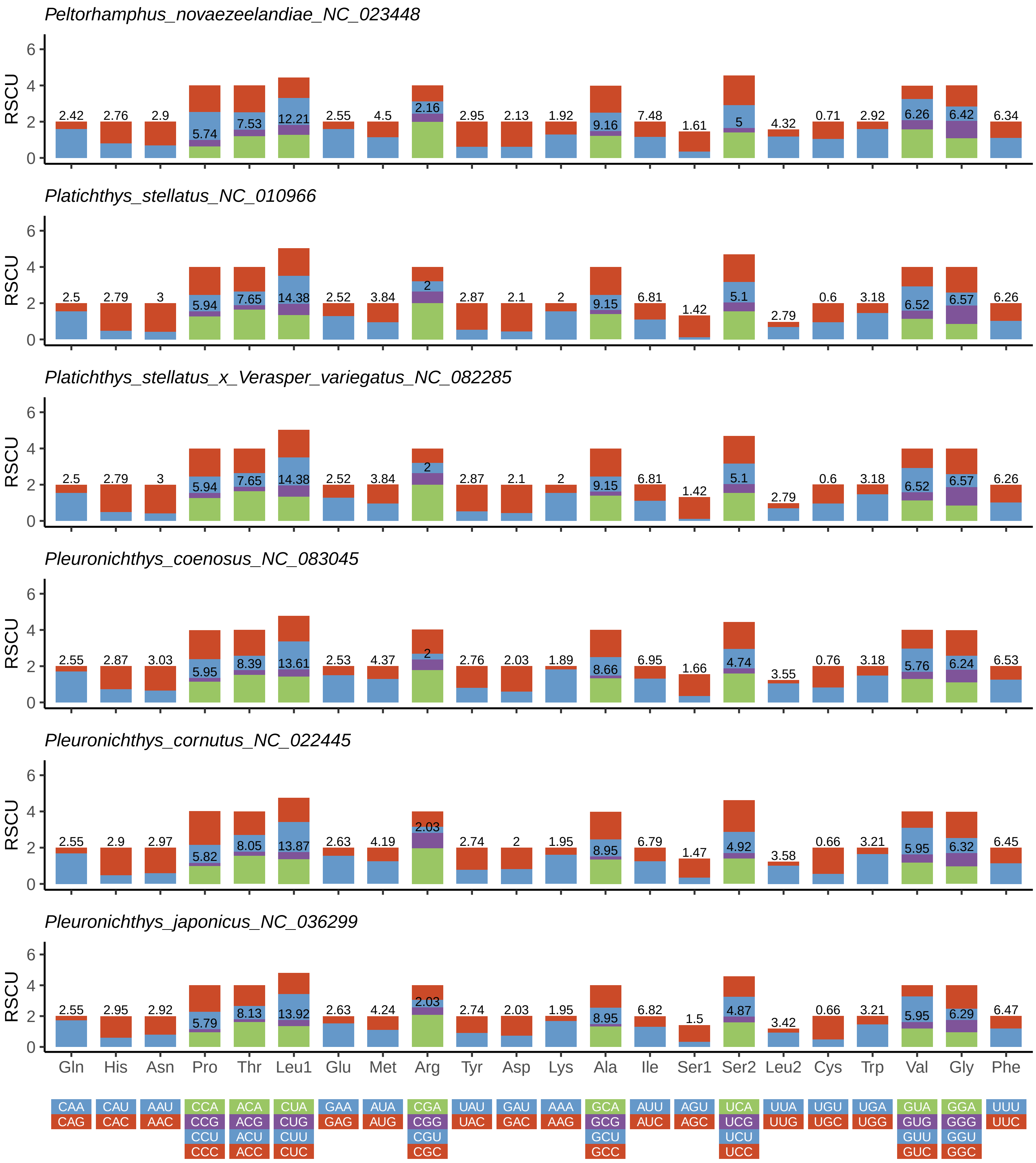
Peltorhamphus_novaezeelandiae_NC_023448
2.42 2.76 2.9 5.74 7.53 12.21 2.55 4.5 2.16 2.95 2.13 1.92 9.16 7.48 1.61 5 4.32 0.71 2.92 6.26 6.42 6.34
Platichthys_stellatus_NC_010966
2.5 2.79 3 5.94 7.65 14.38 2.52 3.84 2 2.87 2.1 2 9.15 6.81 1.42 5.1 2.79 0.6 3.18 6.52 6.57 6.26
Platichthys_stellatus_x_Verasper_variegatus_NC_082285
2.5 2.79 3 5.94 7.65 14.38 2.52 3.84 2 2.87 2.1 2 9.15 6.81 1.42 5.1 2.79 0.6 3.18 6.52 6.57 6.26
Pleuronichthys_coenosus_NC_083045
2.55 2.87 3.03 5.95 8.39 13.61 2.53 4.37 2 2.76 2.03 1.89 8.66 6.95 1.66 4.74 3.55 0.76 3.18 5.76 6.24 6.53
Pleuronichthys_cornutus_NC_022445
2.55 2.9 2.97 5.82 8.05 13.87 2.63 4.19 2.03 2.74 2 1.95 8.95 6.79 1.47 4.92 3.58 0.66 3.21 5.95 6.32 6.45
Pleuronichthys_japonicus_NC_036299
2.55 2.95 2.92 5.79 8.13 13.92 2.63 4.24 2.03 2.74 2.03 1.95 8.95 6.82 1.5 4.87 3.42 0.66 3.21 5.95 6.29 6.47
RSCU
0 2 4 6
Gln His Asn Pro Thr Leu1 Glu Met Arg Tyr Asp Lys Ala Ile Ser1 Ser2 Leu2 Cys Trp Val Gly Phe
CAA CAU AAU CCA ACA CUA GAA AUA CGA UAU GAU AAA GCA AUU AGU UCA UUA UGU UGA GUA GGA UUU
CAG CAC AAC CCG ACG CUG GAG AUG CGG UAC GAC AAG GCG AUC AGC UCG UUG UGC UGG GUG GGG UUC
CCU ACU CUU CGU GCU UCU GUU GGU
CCC ACC CUC CGC GCC UCC GUC GGC

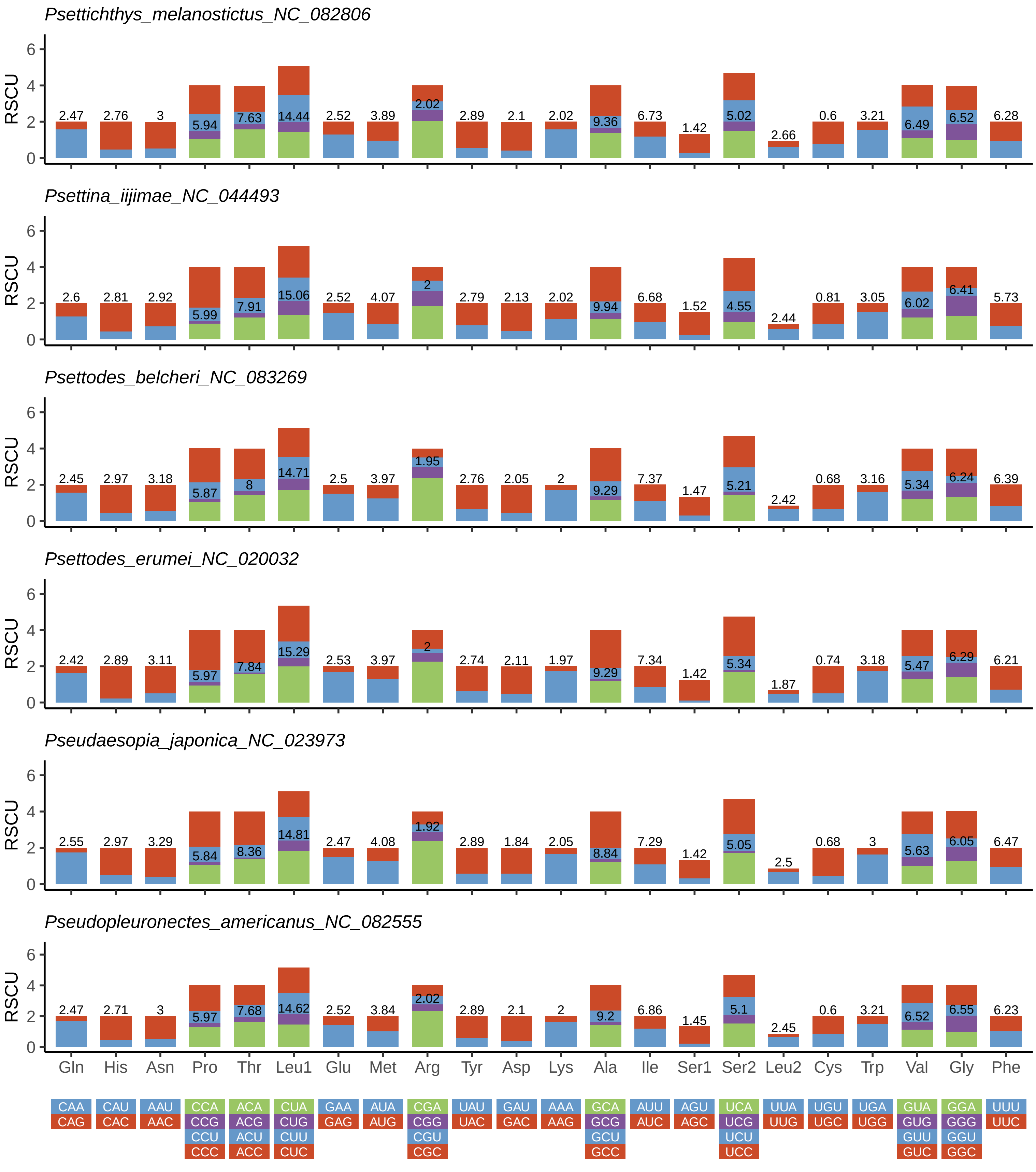
Psettichthys_melanostictus_NC_082806
RSCU
2.47 2.76 3 5.94 7.63 14.44 2.52 3.89 2.02 2.89 2.1 2.02 9.36 6.73 1.42 5.02 2.66 0.6 3.21 6.49 6.52 6.28
Psettina_iijimae_NC_044493
RSCU
2.6 2.81 2.92 5.99 7.91 15.06 2.52 4.07 2 2.79 2.13 2.02 9.94 6.68 1.52 4.55 2.44 0.81 3.05 6.02 6.41 5.73
Psettodes_belcheri_NC_083269
RSCU
2.45 2.97 3.18 5.87 8 14.71 2.5 3.97 1.95 2.76 2.05 2 9.29 7.37 1.47 5.21 2.42 0.68 3.16 5.34 6.24 6.39
Psettodes_erumei_NC_020032
RSCU
2.42 2.89 3.11 5.97 7.84 15.29 2.53 3.97 2 2.74 2.11 1.97 9.29 7.34 1.42 5.34 1.87 0.74 3.18 5.47 6.29 6.21
Pseudaesopia_japonica_NC_023973
RSCU
2.55 2.97 3.29 5.84 8.36 14.81 2.47 4.08 1.92 2.89 1.84 2.05 8.84 7.29 1.42 5.05 2.5 0.68 3 5.63 6.05 6.47
Pseudopleuronectes_americanus_NC_082555
RSCU
2.47 2.71 3 5.97 7.68 14.62 2.52 3.84 2.02 2.89 2.1 2 9.2 6.86 1.45 5.1 2.45 0.6 3.21 6.52 6.55 6.23
Gln His Asn Pro Thr Leu1 Glu Met Arg Tyr Asp Lys Ala Ile Ser1 Ser2 Leu2 Cys Trp Val Gly Phe
CAA CAU AAU CCA ACA CUA GAA AUA CGA UAU GAU AAA GCA AUU AGU UCA UUA UGU UGA GUA GGA UUU
CAG CAC AAC CCG ACG CUG GAG AUG CGG UAC GAC AAG GCG AUC AGC UCG UUG UGC UGG GUG GGG UUC
CCU ACU CUU CGU GCU UCU GUU GGU
CCC ACC CUC CGC GCC UCC GUC GGC

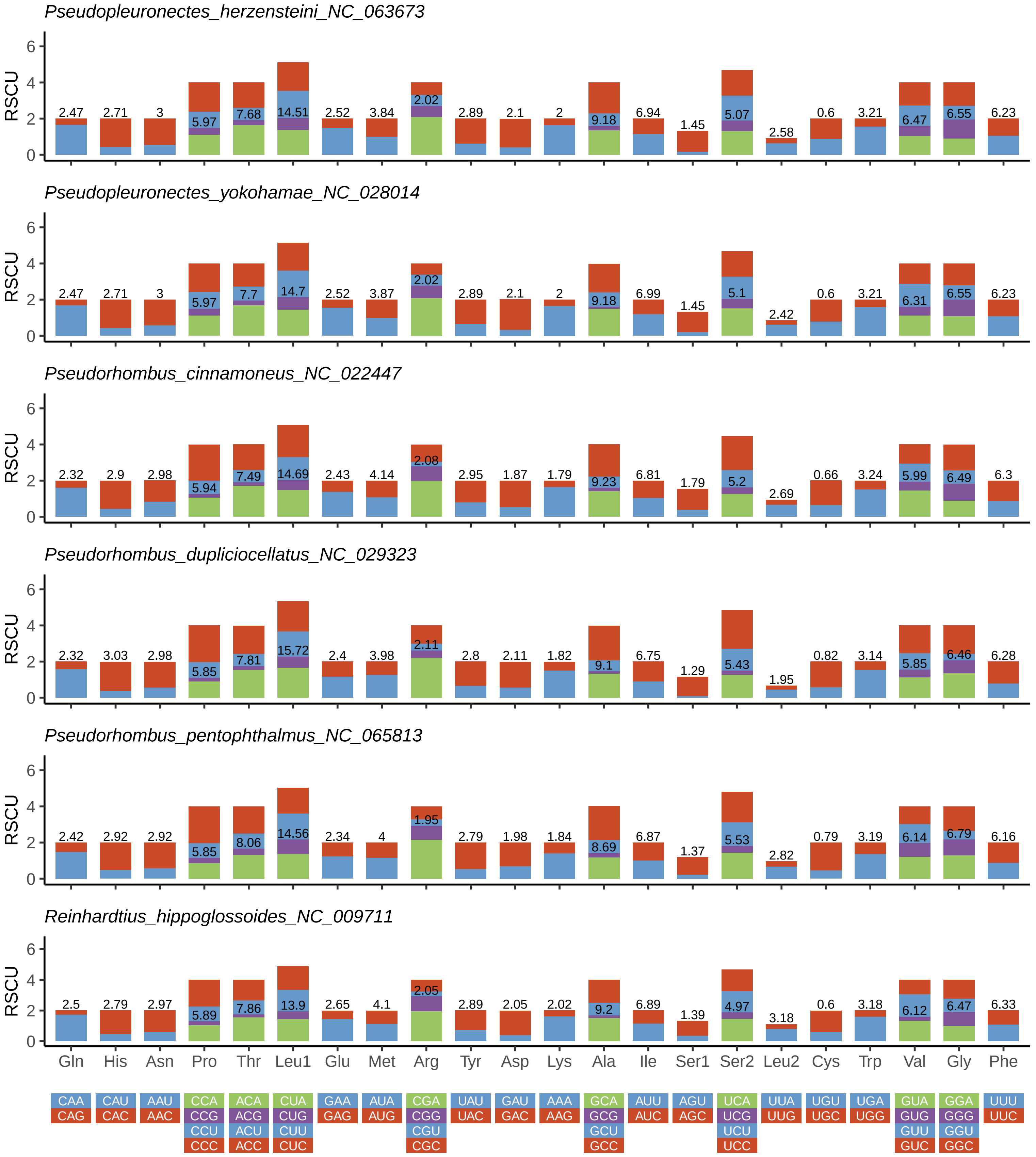
Pseudopleuronectes_herzensteini_NC_063673
2.47 2.71 3 5.97 7.68 14.51 2.52 3.84 2.02 2.89 2.1 2 9.18 6.94 1.45 5.07 2.58 0.6 3.21 6.47 6.55 6.23
Pseudopleuronectes_yokohamae_NC_028014
2.47 2.71 3 5.97 7.7 14.7 2.52 3.87 2.02 2.89 2.1 2 9.18 6.99 1.45 5.1 2.42 0.6 3.21 6.31 6.55 6.23
Pseudorhombus_cinnamoneus_NC_022447
2.32 2.9 2.98 5.94 7.49 14.69 2.43 4.14 2.08 2.95 1.87 1.79 9.23 6.81 1.79 5.2 2.69 0.66 3.24 5.99 6.49 6.3
Pseudorhombus_dupliciocellatus_NC_029323
2.32 3.03 2.98 5.85 7.81 15.72 2.4 3.98 2.11 2.8 2.11 1.82 9.1 6.75 1.29 5.43 1.95 0.82 3.14 5.85 6.46 6.28
Pseudorhombus_pentophthalmus_NC_065813
2.42 2.92 2.92 5.85 8.06 14.56 2.34 4 1.95 2.79 1.98 1.84 8.69 6.87 1.37 5.53 2.82 0.79 3.19 6.14 6.79 6.16
Reinhardtius_hippoglossoides_NC_009711
2.5 2.79 2.97 5.89 7.86 13.9 2.65 4.1 2.05 2.89 2.05 2.02 9.2 6.89 1.39 4.97 3.18 0.6 3.18 6.12 6.47 6.33
RSCU
0 2 4 6
Gln His Asn Pro Thr Leu1 Glu Met Arg Tyr Asp Lys Ala Ile Ser1 Ser2 Leu2 Cys Trp Val Gly Phe
CAA CAU AAU CCA ACA CUA GAA AUA CGA UAU GAU AAA GCA AUU AGU UCA UUA UGU UGA GUA GGA UUU
CAG CAC AAC CCG ACG CUG GAG AUG CGG UAC GAC AAG GCG AUC AGC UCG UUG UGC UGG GUG GGG UUC
CCU ACU CUU CGU GCU UCU GUU GGU
CCC ACC CUC CGC GCC UCC GUC GGC

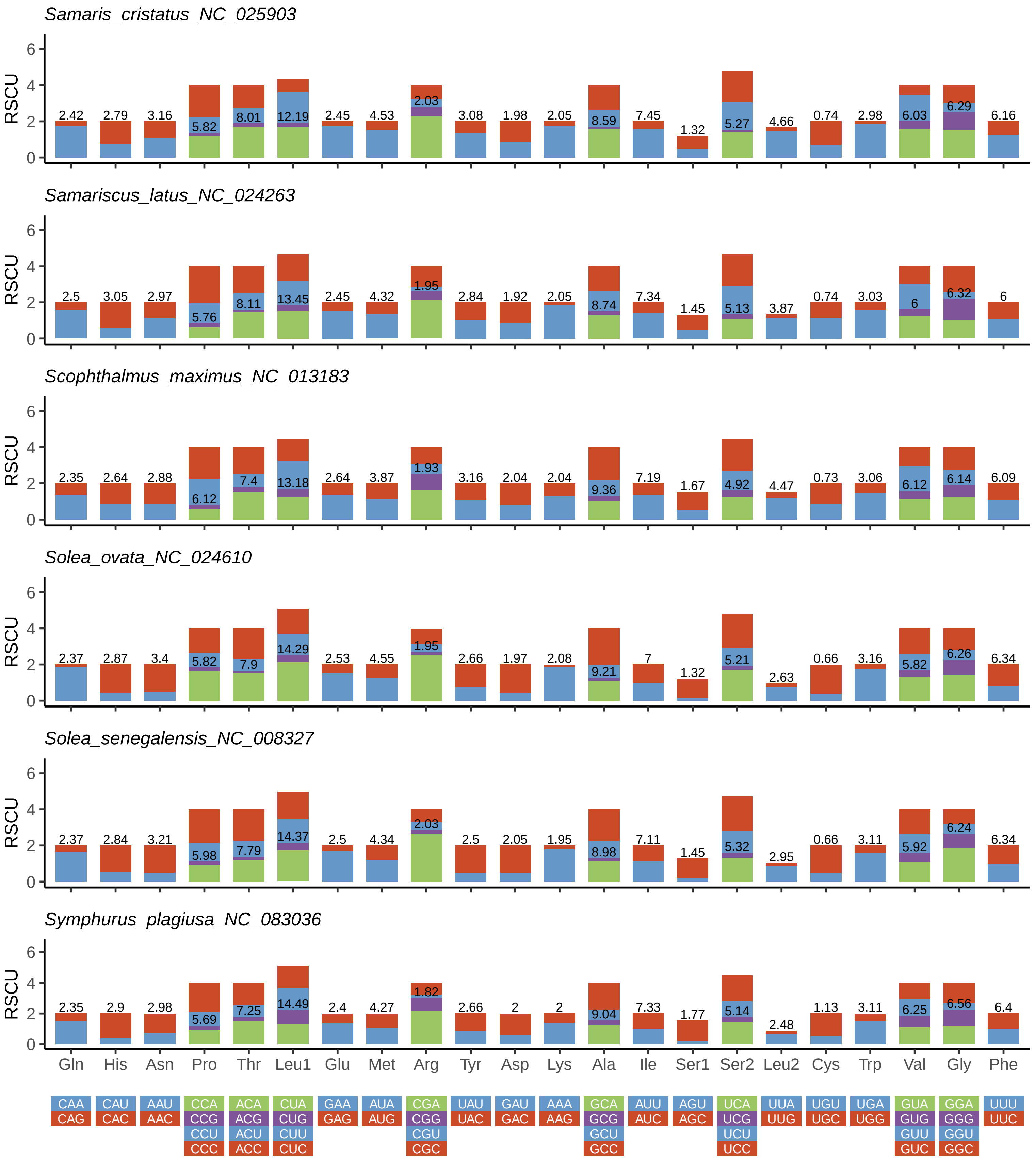
Samaris_cristatus_NC_025903
Samariscus_latus_NC_024263
Scophthalmus_maximus_NC_013183
Solea_ovata_NC_024610
Solea_senegalensis_NC_008327
Symphurus_plagiusa_NC_083036
RSCU
Gln His Asn Pro Thr Leu1 Glu Met Arg Tyr Asp Lys Ala Ile Ser1 Ser2 Leu2 Cys Trp Val Gly Phe
CAA CAU AAU CCA ACA CUA GAA AUA CGA UAU GAU AAA GCA AUU AGU UCA UUA UGU UGA GUA GGA UUU
CAG CAC AAC CCG ACG CUG GAG AUG CGG UAC GAC AAG GCG AUC AGC UCG UUG UGC UGG GUG GGG UUC
CCU ACU CUU CGU GCU UCU GUU GGU
CCC ACC CUC CGC GCC UCC GUC GGC

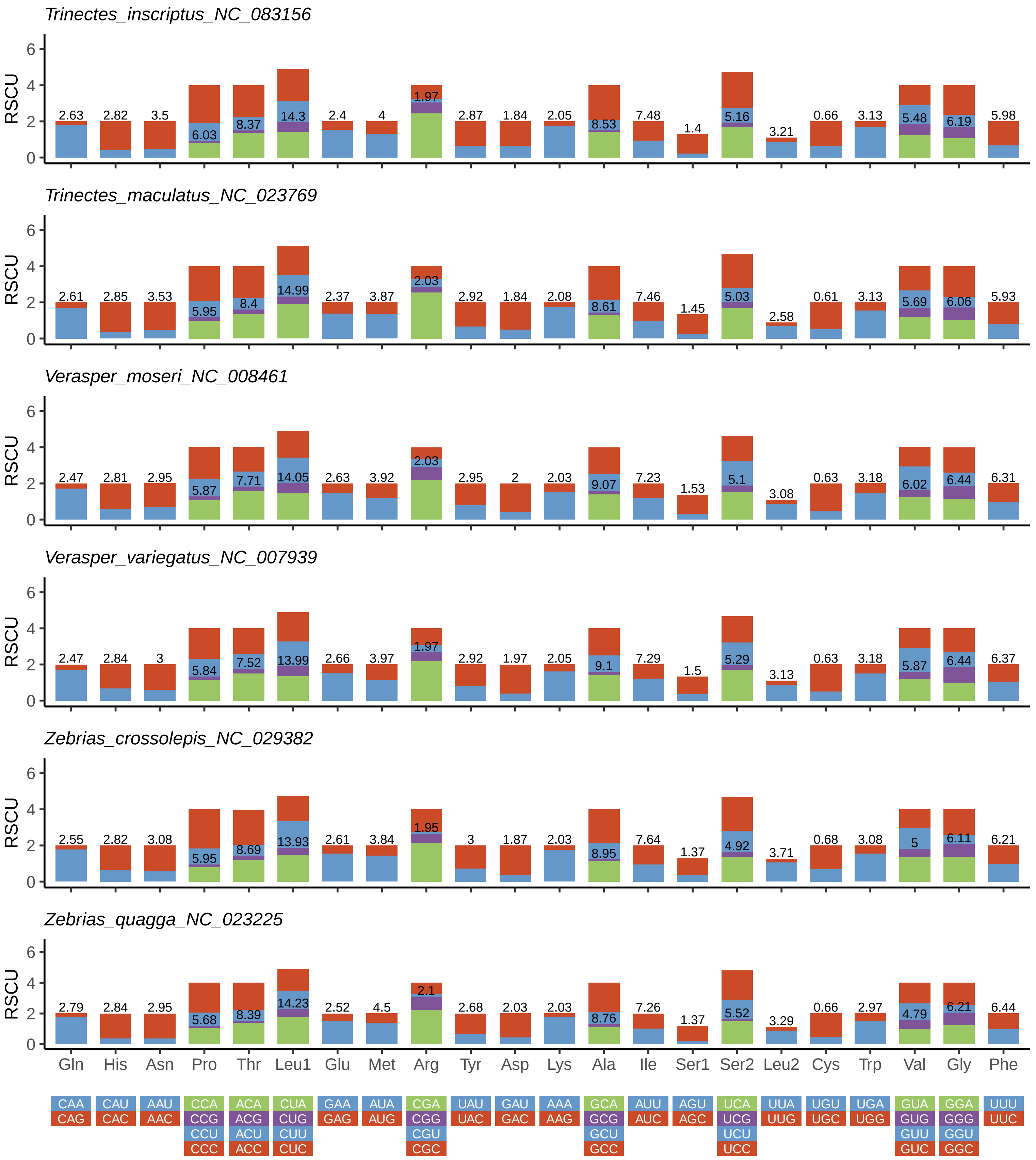
Trinectes_inscriptus_NC_083156
RSCU
2.63 2.82 3.5 6.03 8.37 14.3 2.4 4 1.97 2.87 1.84 2.05 8.53 7.48 1.4 5.16 3.21 0.66 3.13 5.48 6.19 5.98
Trinectes_maculatus_NC_023769
RSCU
2.61 2.85 3.53 5.95 8.4 14.99 2.37 3.87 2.03 2.92 1.84 2.08 8.61 7.46 1.45 5.03 2.58 0.61 3.13 5.69 6.06 5.93
Verasper_moseri_NC_008461
RSCU
2.47 2.81 2.95 5.87 7.71 14.05 2.63 3.92 2.03 2.95 2 2.03 9.07 7.23 1.53 5.1 3.08 0.63 3.18 6.02 6.44 6.31
Verasper_variegatus_NC_007939
RSCU
2.47 2.84 3 5.84 7.52 13.99 2.66 3.97 1.97 2.92 1.97 2.05 9.1 7.29 1.5 5.29 3.13 0.63 3.18 5.87 6.44 6.37
Zebrias_crossolepis_NC_029382
RSCU
2.55 2.82 3.08 5.95 8.69 13.93 2.61 3.84 1.95 3 1.87 2.03 8.95 7.64 1.37 4.92 3.71 0.68 3.08 5 6.11 6.21
Zebrias_quagga_NC_023225
RSCU
2.79 2.84 2.95 5.68 8.39 14.23 2.52 4.5 2.1 2.68 2.03 2.03 8.76 7.26 1.37 5.52 3.29 0.66 2.97 4.79 6.21 6.44
0 2 4 6
Gln His Asn Pro Thr Leu1 Glu Met Arg Tyr Asp Lys Ala Ile Ser1 Ser2 Leu2 Cys Trp Val Gly Phe
CAA CAG
CAU CAC
AAU AAC
CCA CCG CCU CCC
ACA ACG ACU ACC
CUA CUG CUU CUC
GAA GAG
AUA AUG
CGA CGG CGU CGC
UAU UAC
GAU GAC
AAA AAG
GCA GCG GCU GCC
AUU AUC
AGU AGC
UCA UCG UCU UCC
UUA UUG
UGU UGC
UGA UGG
GUA GUG GUU GUC
GGA GGG GGU GGC
UUU UUC

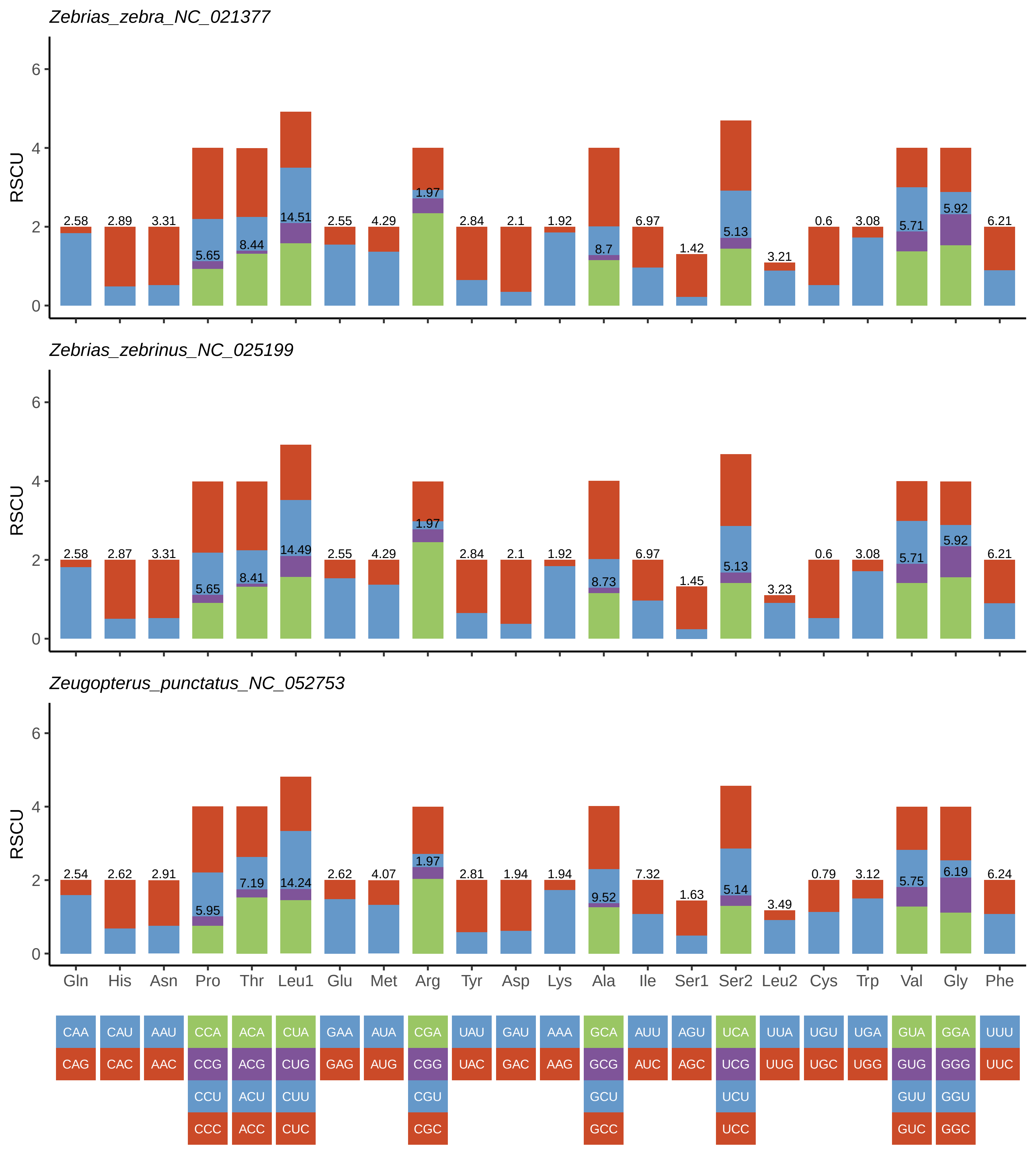
Zebrias_zebra_NC_021377
RSCU
0 2 4 6
2.58 2.89 3.31 5.65 8.44 14.51 2.55 4.29 1.97 2.84 2.1 1.92 8.7 6.97 1.42 5.13 3.21 0.6 3.08 5.71 5.92 6.21
Zebrias_zebrinus_NC_025199
RSCU
0 2 4 6
2.58 2.87 3.31 5.65 8.41 14.49 2.55 4.29 1.97 2.84 2.1 1.92 8.73 6.97 1.45 5.13 3.23 0.6 3.08 5.71 5.92 6.21
Zeugopterus_punctatus_NC_052753
RSCU
0 2 4 6
2.54 2.62 2.91 5.95 7.19 14.24 2.62 4.07 1.97 2.81 1.94 1.94 9.52 7.32 1.63 5.14 3.49 0.79 3.12 5.75 6.19 6.24
Gln His Asn Pro Thr Leu1 Glu Met Arg Tyr Asp Lys Ala Ile Ser1 Ser2 Leu2 Cys Trp Val Gly Phe
CAA CAU AAU CCA ACA CUA GAA AUA CGA UAU GAU AAA GCA AUU AGU UCA UUA UGU UGA GUA GGA UUU
CAG CAC AAC CCG ACG CUG GAG AUG CGG UAC GAC AAG GCG AUC AGC UCG UUG UGC UGG GUG GGG UUC
CCU ACU CUU CGU GCU UCU GUU GGU
CCC ACC CUC CGC GCC UCC GUC GGC

**Fig. S3.** Relative synonymous codon usage of protein-coding genes in mitochondrial genomes of studied flatfish.
