## Supplementary material for "Comprehensive analysis of 111 Pleuronectiformes mitochondrial genomes: insights into structure, conservation, variation and evolution": Fig. S1

**A**

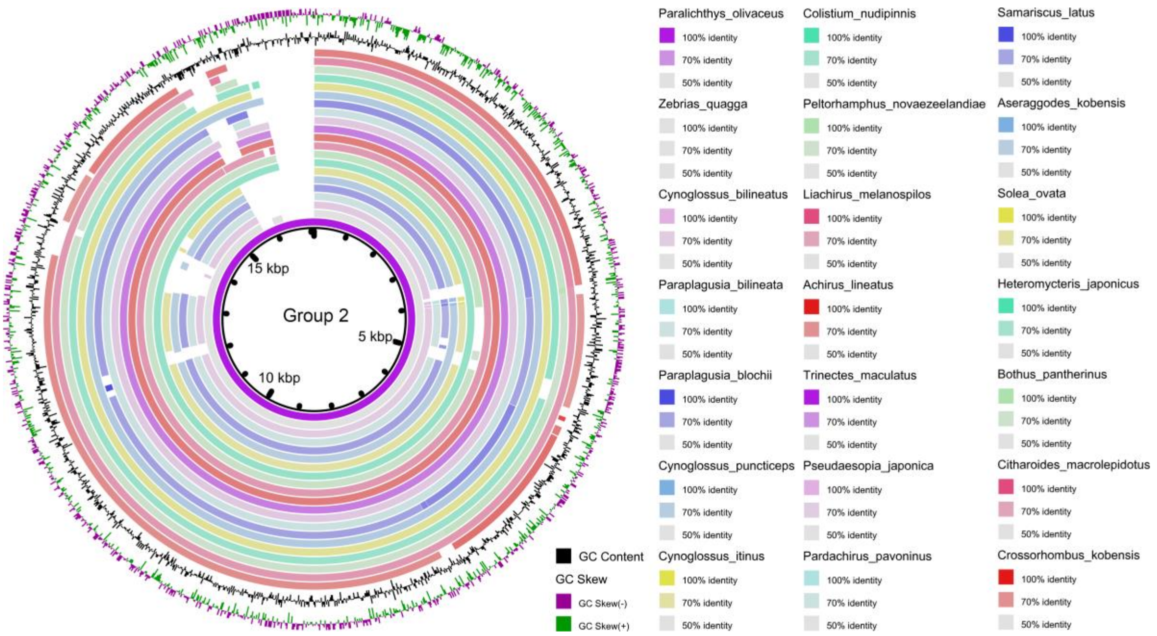
Group 2
15 kbp
5 kbp
10 kbp
Paralichthys_olivaceus
Colistium_nudipinnis
Samariscus_latus
Zebrias_quagga
Peltorhamphus_novaezeelandiae
Aseraggodes_kobensis
Cynoglossus_bilineatus
Liachirus_melanospilos
Solea_ovata
Paraplagusia_bilineata
Achirus_lineatus
Heteromycteris_japonicus
Paraplagusia_blochii
Trinectes_maculatus
Bothus_pantherinus
Cynoglossus_puncticeps
Pseudaesopia_japonica
Citharoides_macrolepidotus
Cynoglossus_itinus
Pardachirus_pavoninus
Crossorhombus_kobensis
100% identity
70% identity
50% identity
GC Content
GC Skew
GC Skew(-)
GC Skew(+)

**B**

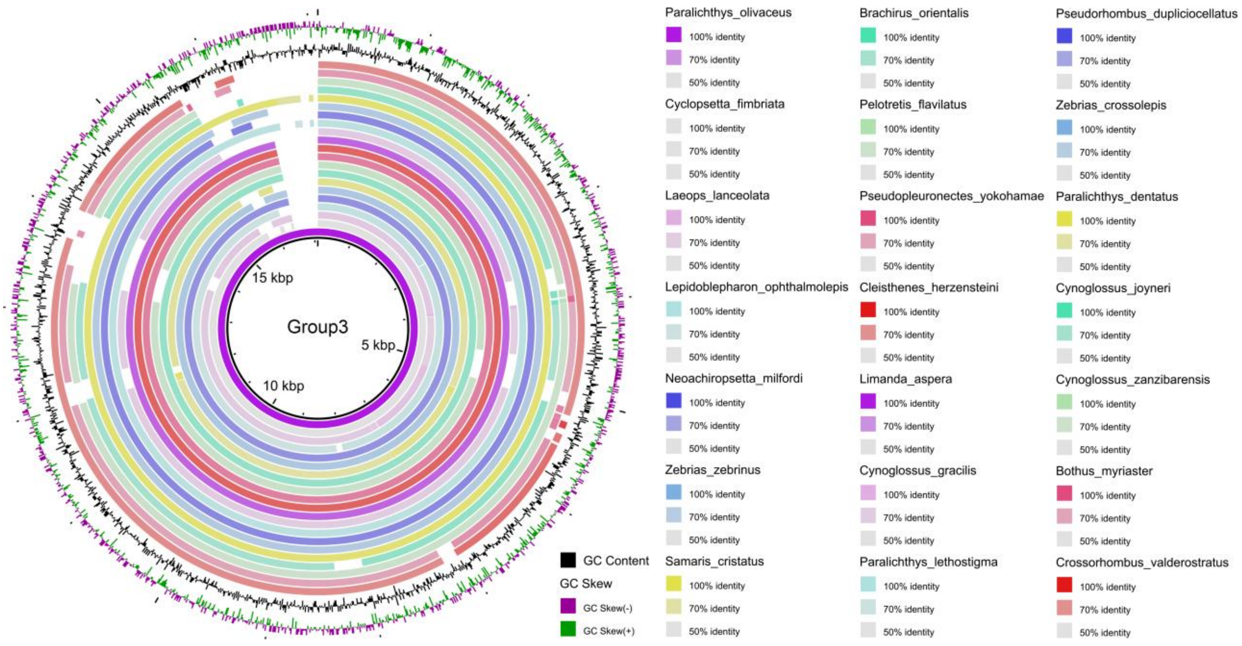
Group3
15 kbp
5 kbp
10 kbp
Paralichthys_olivaceus
Brachirus_orientalis
Pseudorhombus_dupliciocellatus
Cyclopsetta_fimbriata
Pelotretis_flavilatus
Zebrias_crossolepis
Laeops_lanceolata
Pseudopleuronectes_yokohamae
Paralichthys_dentatus
Lepidoblepharon_ophthalmolepis
Cleisthenes_herzensteini
Cynoglossus_joyneri
Neoachiropsetta_milfordi
Limanda_aspera
Cynoglossus_zanzibarensis
Zebrias_zebrinus
Cynoglossus_gracilis
Bothus_myriaster
Samaris_cristatus
Paralichthys_lethostigma
Crossorhombus_valderostratus
100% identity
70% identity
50% identity
GC Content
GC Skew
GC Skew(-)
GC Skew(+)

**C**

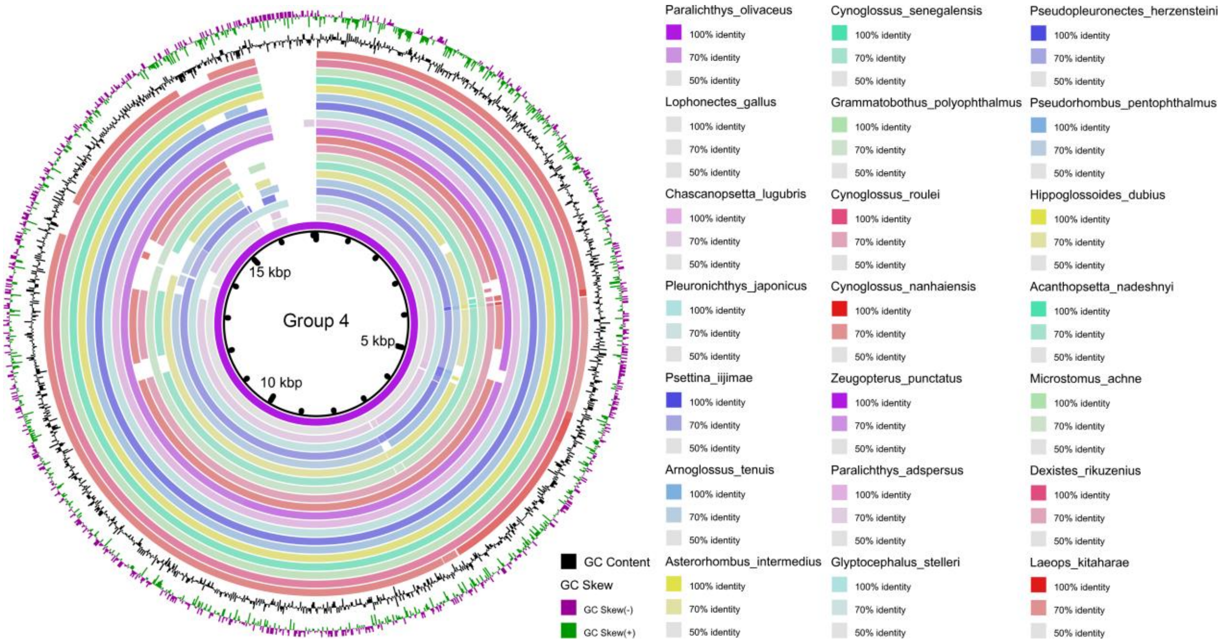

15 kbp
Group 4
5 kbp
10 kbp
Paralichthys_olivaceus
Cynoglossus_senegalensis
Pseudopleuronectes_herzensteini
Lophonectes_gallus
Grammatobothus_polyophthalmus
Pseudorhombus_pentophthalmus
Chascanopsetta_lugubris
Cynoglossus_roulei
Hippoglossoides_dubius
Pleuronichthys_japonicus
Cynoglossus_nanhaiensis
Acanthopsetta_nadeshnyi
Psettina_iijimae
Zeugopterus_punctatus
Microstomus_achne
Arnoglossus_tenuis
Paralichthys_adspersus
Dexistes_rikuzenius
Asterorhombus_intermedius
Glyptocephalus_stelleri
Laeops_kitaharae
100% identity
70% identity
50% identity
GC Content
GC Skew
GC Skew(-)
GC Skew(+)


**D**

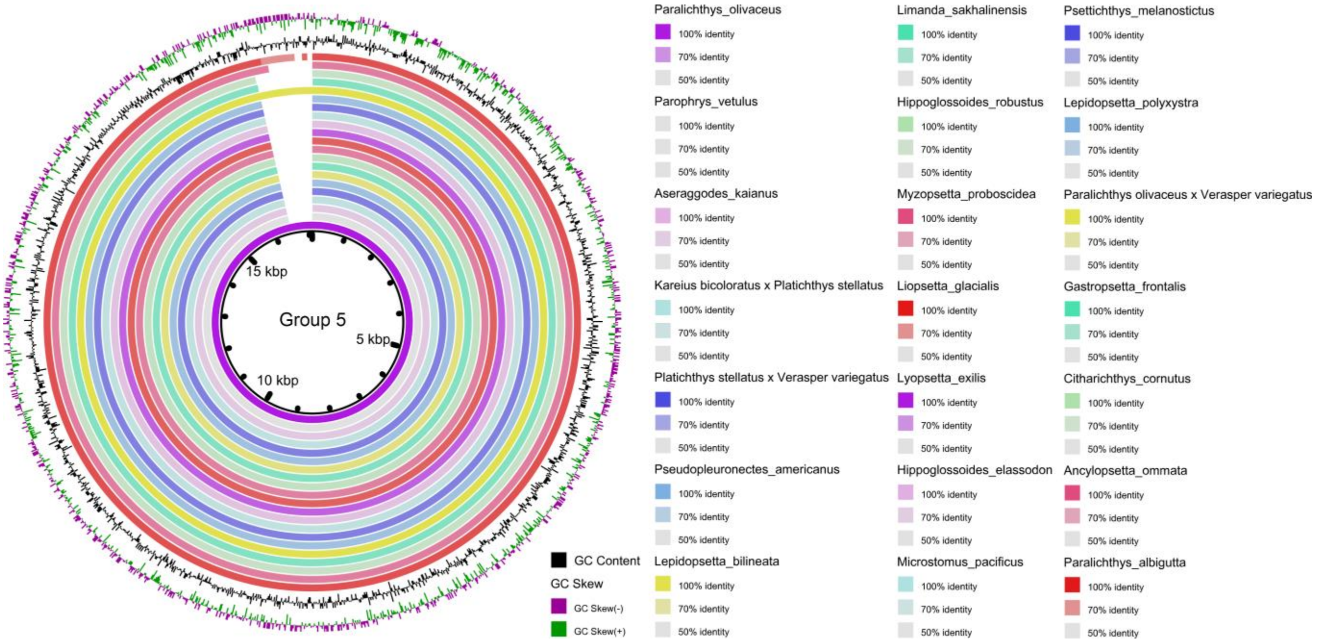

15 kbp
Group 5
5 kbp
10 kbp
Paralichthys_olivaceus
Limanda_sakhalinensis
Psettichthys_melanostictus
Parophrys_vetulus
Hippoglossoides_robustus
Lepidopsetta_polyxystra
Aseraggodes_kaianus
Myzopsetta_proboscidea
Paralichthys olivaceus x Verasper variegatus
Kareius bicoloratus x Platichthys stellatus
Liopsetta_glacialis
Gastropsetta_frontalis
Platichthys stellatus x Verasper variegatus
Lyopsetta_exilis
Citharichthys_cornutus
Pseudopleuronectes_americanus
Hippoglossoides_elassodon
Ancylopsetta_ommata
Lepidopsetta_bilineata
Microstomus_pacificus
Paralichthys_albigutta
100% identity
70% identity
50% identity
GC Content
GC Skew
GC Skew(-)
GC Skew(+)

**E**

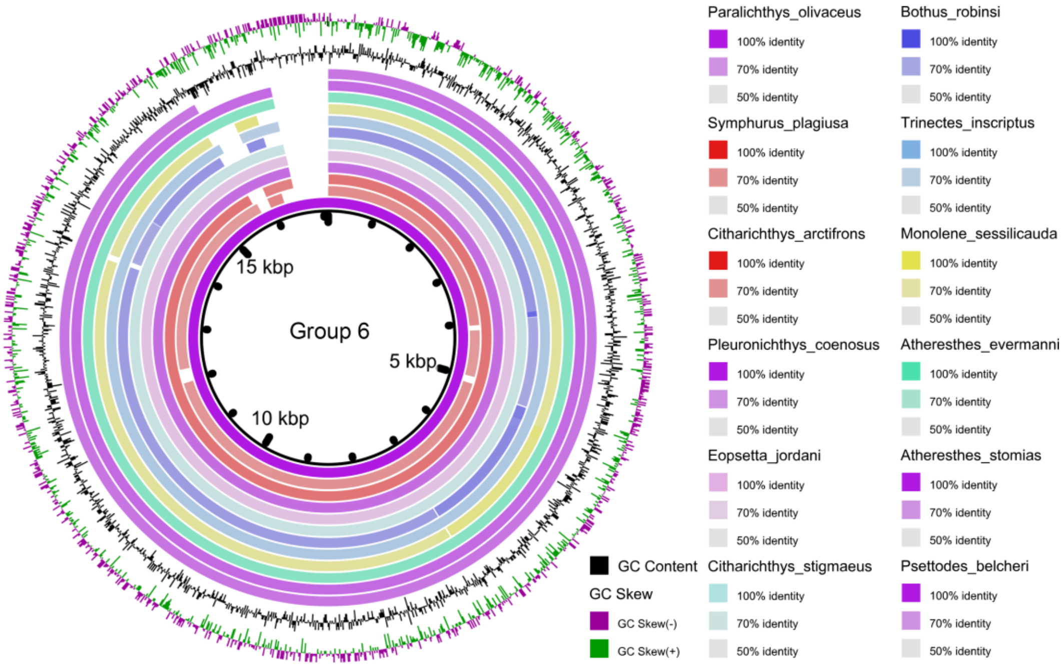


Fig. S1. Alignment of mitochondrial genomes among Pleuronectiformes species (Group 2-6). The gap in the circle represents mismatched sequence of genome alignment. GC content (black) and GC skew (purple/green) are at the outermost two circles. Increased GC content and positive GC skew are represented by peaks oriented toward the center of the circle.
