## Supplementary material for "Comprehensive analysis of 111 Pleuronectiformes mitochondrial genomes: insights into structure, conservation, variation and evolution": Fig. S2

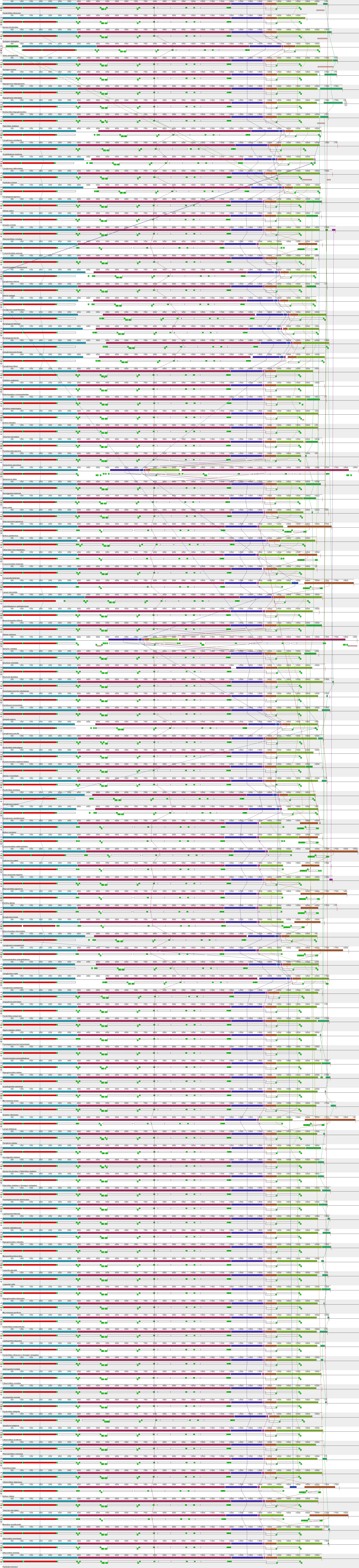

**Fig. S2.** Collinearity analysis of mitochondrial genomes of studied flatfish. For each species, the first row represents the collinear blocks, and the second row represents the genes, with rRNA genes indicated in red, protein-coding genes in white, tRNA genes in green, and repeat region in pink. co-linear blocks are connected by lines of the same color.
